# Proteomic Changes in the Cytoplasmatic Fraction of Weaned Piglets Liver and Kidney Under Antioxidants and Mycotoxins Diets

**DOI:** 10.1101/2024.05.13.593823

**Authors:** Roua Gabriela Popescu, Anca Dinischiotu, George Cătălin Marinescu

**Affiliations:** Independent Research Association, Bucharest 012416, Romania; Blue Screen SRL, Bucharest 012416, Romania; Department of Biochemistry and Molecular Biology, Faculty of Biology, University of Bucharest, Bucharest 050095, Romania

**Keywords:** piglets, mycotoxins, proteomics, DIA, feed additives, antioxidant effect, antioxidants proteomics, cytoplasmatic fraction, pathway analysis

## Abstract

Mycotoxin contamination represents a threat for human and animal health. Antioxidants can mitigate some of these effects due to their ability to scavenge free radicals, reduce oxidative stress, and have anti-inflammatory and immunomodulatory properties. This work investigates the potential of antioxidants derived from grape seed and sea buckthorn to mitigate the adverse effects of aflatoxin B1 (AFB1) and ochratoxin A (OTA), as mycotoxins contamination in weaned piglet diets. We used an unbiased Data Independent Acquisition (DIA) approach to analyse the proteomic impact of OTA and AFB1-contaminated diets on liver and kidney cytoplasmic metabolism, particularly focusing on the conjugation phase. We show that some of the mycotoxins effects are partial mitigated by the antioxidants enriched diet. Additionally, in kidneys, some of the effects are synergistically amplified, such as proteins involved in the fatty acids degradation, peroxisome, PPAR signalling, translation, TCA cycle and the excretion pathways. Inclusion of antioxidants in the animal diet can have beneficial effects. Nevertheless, caution is advised, synergistic effects can occur with potentially more serious consequences than the effect of mycotoxins alone.

## 1. Introduction

Certain strains of fungi growing on food and feed crops produce small and stable toxic secondary metabolites known as mycotoxins, which occur in various stages of food production (Haque et al. 2020; Pandey et al. 2023). Mycotoxin contamination represents a threat for human and animal health, especially through the negative effects on hormonal, immune and DNA repair functions as well as lipid, carbohydrate, and amino acid metabolism (Omotayo et al. 2019; Malvandi et al. 2022). Mycotoxins enter the body through consumption of contaminated processed food or its primary constituents, such as cereals, oilseeds, fruits, and vegetables, either directly or as by-products (Kepinska-Pacelik and Biel 2021; Abdolmaleki et al., 2021). In addition, humans are more exposed to mycotoxin contamination through the consumption of milk, eggs, meat, and fish derived from animals that were fed a mycotoxin-contaminated diet (Carballo et al., 2018; Adegbeye et al., 2020). Following absorption through the digestive system, mycotoxins are distributed in different organs of the body and metabolized, mainly by the liver and kidney (Tolosa et al. 2021).

Once up-taken in the body, mycotoxins are metabolised by the biotransformation of xenobiotics pathway. In phase I, namely the modification phase, members of the cytochromes P450, which mainly catalyse the oxygenation reactions, as well as the dealkylation reactions and those for the formation of epoxides (Dang et al. 2020). In phase II or conjugation phase, glutathione S-transferases, sulfotransferases, N-acetyltransferases, and uridine 5’-diphospho-(UDP)-glucuronosyltransferases are involved (Jancova et al. 2010). Finally, in phase III, respectively the excretion phase, transporters present in the liver, kidneys and intestines facilitate the elimination of metabolites formed in cells in an active manner (Lehman-McKeeman and Ruepp 2018; Li et al. 2019).

Currently, numerous strategies have been developed to prevent, reduce, or even eliminate mycotoxin contamination in animal feed through biological, chemical, and physical detoxification methods, which allow the degradation of mycotoxins and their corresponding metabolites and maintain the nutritional value of food products, without introducing other potentially toxic substances into biological systems (Luo et al. 2018; Čolović et al. 2019; Hamad et al. 2023). Among these, the most promising method is the addition of antioxidants derived from plants to feed to reduce the harmful effects caused by the presence of mycotoxins on animal health. Several studies suggest that antioxidants can counteract chemical carcinogenesis by being administered before or concurrently with the carcinogen, thereby supporting their inhibitory potential against the effects of mycotoxins (Mut-Salud et al. 2016; Van Le Thanh et al. 2016; Deng et al. 2019). Although adding antioxidants to feed could support the circular economy (Arias et al. 2022), it is usually not feasible due to various factors. In addition, not all antioxidants are suitable for all types of feed due to the problems of color, taste, solubility, stability, biological activity, and interaction with other feed components (like proteins or phenolic compounds) (Hadidi et al. 2022).

An interesting fact is that in certain cases the sources of potential protective agents are also a source of mycotoxins, especially the most common AFB1 and OTA. This scenario can be encountered with coffee, tea, citrus and herbs (Abdel-Wahhab and Kholif 2008; Chalyy et al. 2021). Therefore, the prevention of effects should strictly promote anti-mycotoxin action. To date, most studies have tried to confirm the beneficial effects of adding food by-products rich in antioxidants to the animal diet. However, in recent years, with the development of proteomics and metabolomics methods, new studies have highlighted their side or even negative effects (Petcu et al. 2023; Popescu et al. 2023). Antioxidants are believed to counteract some of the effects of mycotoxins due to their ability to scavenge free radicals, reduce oxidative stress, and have anti-inflammatory and immunomodulatory properties (Chaudhary et al. 2023). The specific mechanisms by which antioxidants mitigate or counteract the effects of mycotoxins may vary depending on the type of antioxidant and the mycotoxin involved.

In our previous investigation about the effects of feed mycotoxin contamination with AFB1 and OTA, and the potential of addition of by-product mixture of antioxidants to mitigate these effects on microsomal fraction of weaned piglets’ liver, we found that some proteins expressions have been affected by antioxidants (Popescu et al. 2023). However, these observations only apply to the microsomal fraction that contains mostly proteins involved in phase I biotransformation, while phase II or conjugation proteins are located within the cytoplasmatic fraction.

In this study, we aimed to investigate the proteomic level impact of OTA and AFB1 contaminated-diet in piglets using an unbiased Data Independent Acquisition (DIA) approach, showing how the addition of grape seed and sea buckthorn meal to the diet impacts liver and kidney cytoplasmatic metabolism in conjugation phase.

## 2. Material and methods

### 2.1. Animal experiments

For a 30-day feeding trial, a total of 40 weaned piglets of the TOPIGS-40 hybrid breed (average body weight of 9.11 kg±0.03 kg) were divided into four groups. The control group (C) was fed with a standard diet for starter piglets. Experimental groups included: antioxidants group (A), which received the basal diet supplemented with a 5% mixture of grape seed and sea buckthorn meal by-products; mycotoxins group (M), received the basal diet contaminated with 62 ppb of aflatoxin B1 (AFB1) and 479 ppb of ochratoxin A (OTA); and antioxidants plus mycotoxins co-administration group (AM), received the basal diet with by-products mixture and contaminated with AFB1 and OTA. Diets were formulated according to Popescu et al. (2021a). Animal performance and biomarkers of liver and kidney function have been previously reported (Popescu et al. 2021b). All animals had *ad libitum* access to water and assigned diets throughout the study. The experiment took place at the National Research-Development Institute for Animal Nutrition and Biology, Balotești, Romania. The Ethical Committee of it approved all experimental procedures regarding animals (*n =* 4 per group) at the end of the 30-day period, in compliance with EU Council Directive 98/58/EC and Romanian Law 206/2004 (Ethical Committee no. 118/02.12.2019). Samples of liver and kidney were collected and stored at - 80°C.

### 2.2. Isolation of the cytoplasmatic fraction

The cytoplasmatic fraction isolation method was conducted with modifications of the protocol described by Rasmussen et al. (2011). Six grams of sample tissue were minced and homogenized in ice-cold Tris-sucrose buffer using a Potter homogenizer. The resulted homogenate was centrifuged at 10,000 x g for 10 minutes at 4 °C to obtain the crude homogenate without nuclei. This obtained crude homogenate was then centrifuged at 100,000 *g* for 60 minutes at 4 °C, resulting the cytoplasmatic fraction in the supernatant, and the microsomal fraction in the pellet. The cytoplasmatic supernatant was collected and stored at -80 °C. Purity assay was conducted via immunoblotting (as described previously by Popescu et al. 2021b). All procedures were carried out on ice.

### 2.3. Protein digestion and LC-MS analysis

The cytoplasmatic fraction obtained was subjected to a proteomic analysis. The Bradford method was used to determine the protein concentration. A quantity of 30 μg of protein was resuspended in 500 μL of 50 mM NH_4_HCO_3_, treated with 25 μL of 100 mM DTT at 37 °C for 45 minutes, alkylated by adding 26.25 μL of 300 mM IAA and incubated for another 45 minutes at 37 °C in the dark. Trypsin digestion was performed with a 1 μg /μL trypsin solution (Trypsin Gold, V528A, Promega) at a ratio of 1:50 at 37 °C for 16 h. Digestion was terminated by adding 10 μL of 10% trifluoroacetic acid.

The peptides were dried using speed-vacuum, resuspended in 30 μL of 2% acetonitrile with 0.1% formic acid, and fractionated using a NanoLC 425 system (Eksigent) in a trap-elute configuration. This system included a trapping column C18 (5 μm, 300 μM ID, 25 mm length) and an analytical column Eksigent 5C18-CL-120 (300 μM ID, 150 mm length) connected to DuoSpray ion source (AB Sciex). A volume of 5 μL of peptides were loaded onto trap column at a flow rate of 40 μL /min with 0.1% formic acid and eluted with a gradient from 5 to 80% acetonitrile with 0.1% formic acid over 105 min at a flow rate of 5 μL /min at 55 °C. Ionization was done using electro spray ionization in positive ion mode at a voltage of 5500 V and 200 °C source temperature. The TRIPLE TOF 5600+ operated in Data Independent Acquisition (DIA) mode, with 64 variable windows (SWATH 64 vw) as described by Popescu et al. (2023). The mass spectrometry proteomics data have been deposited to the ProteomeXchange Consortium via the PRIDE (Perez-Riverol et al. 2022) partner repository with the dataset identifier PXD050835 (*Reviewer account details:* **Username:**; **Password:** YD4S7bda). All samples in this study were run in triplicate.

### 2.4. Data analysis

Label-free, library-free protein identification from DIA data was conducted using DIA-NN ver. 1.8.1 (Demichev et al. 2020), searched the raw spectra against the fasta file of the complete *Sus scrofa proteome (*UniProt, UP000008227, January 2024, 46 177) with a precursor m/z range of 400–1250 and trypsin as digestion enzyme. The data were searched with match between runs (MBR) enabled, robust LC quantification, retention time dependent normalisation, and false discovery rate (FDR) < 0.01. Cysteine carbamidomethylation (fixed) and methionine oxidation (variable) modifications were selected. Quantitation was performed using MaxLFQ algorithm. The unique gene matrix *tsv file from DIA-NN was utilized for performing statistical and differential downstream analysis using Limma test with locally installed PolySTest version 1.3 (Schwämmle et al. 2020), available at: https://bitbucket.org/veitveit/polystest/src/master/. For all data, FDR adjusted *p-*value threshold was set to 0.01 and log_2_ fold change (log_2_FC) outside of (-1, 1) interval to highlight the most significantly down- and up-regulated proteins. Following statistical analysis, expression profiles, heatmaps, and plots were performed using the R studio platform (version 2023.12.1+402) with ‘org.Ssc.eg.db’ R package in Bioconductor for *Sus scrofa* (Carlson 2019). Differential pathways expression analysis (PEA) was analysed with PathfindR package, version 1.6.4 (Ulgen et al. 2019), using a protein-protein interaction network (PIN) analysis adapted approach, with PIN data for *Sus scrofa* from STRING (https://stringdb-static.org/download/protein.links.v11.5/). For pathway enrichment analysis (PEA), enrichment chart and term graph, which represented pathways sorted by lowest *p-*value, were generated. Finally, PathfindR output was utilized for integrating and visualising of the data for the main biological pathways using Pathview R package version 1.31.1, (Luo and Brouwer, 2013) and the KEGG pathway database (https://www.genome.jp/kegg/pathway.html/).

## 3. Results

In a previous study (Popescu et al. 2023), the microsomal fraction extracted from weaned piglets exposed to a mycotoxin-contaminated diet, along with an antioxidant additive diet was investigated, to highlight the impact level of AFB1 and OTA on the expression of specific proteins involved in biotransformation metabolism. Since xenobiotic biotransformation involves not only microsomal but also cytoplasmatic proteins related to glycolysis, amino acid metabolism, glycogen metabolism, fatty acid synthesis, and others, we extended our research to include the cytoplasmatic fraction extracted from liver and kidney tissues.

All DIA data were analysed in library-free mode using DIA-NN (version 1.8.1) against the *Sus scrofa* proteome fasta file. We have identified a total of 1279 unique proteins and 56023 precursors within liver cytoplasmatic samples, and 1634 unique proteins and 63344 precursors within kidney cytoplasmatic samples, at a false discovery rate (FDR) below 0.01. Detailed results of identified proteins (*p <* 0.05) are provided in the Supplementary Material (**Tables S1** and **S2**). The relative expression of each protein from each experimental group (A, M, and AM) compared to the control group (C) is shown on KEGG pathways created by the Pathview R module (**Document S1**).

### 3.1. Mycotoxins-contaminated diet impair the detoxification capacity of the liver

Comparing the antioxidant feed diet group (A) to the control group (C), we observed distinct alterations of 11 proteins (**Figure 1**A). Based on the lowest FDR adjusted *p*-value, acyl-CoA oxidase 2 (ACOX2) exhibited a slight decreased expression (log2FC = -0.67), followed by phytanoyl-CoA dioxygenase domain containing 1 (PHYHD1) with a more substantial decrease (log2FC = -0.75). On the other hand, only two proteins demonstrated increased expression, ethyl-malonyl-CoA decarboxylase 1 (ECHDC1) (log2FC = 0.87), along with ferritin light chain (FTL) (log2FC = 0.85) (**Figure 1**D). Additionally, hydroxysteroid dehydrogenase like 2 (HSDL2) showed decreased expression (log2FC = -0.52), while glutathione S-transferase kappa 1 (GSTK1) revealed a similar trend (log2FC = -0.58). Other proteins (**Figure S1**), including kynureninase (KYNU), tryptophanyl-tRNA synthetase 1 (WARS1), and clathrin heavy chain (CLTC), also indicated varying degrees of differential expression, suggesting potential regulatory roles in response to the antioxidant diet.

**Figure 1.**
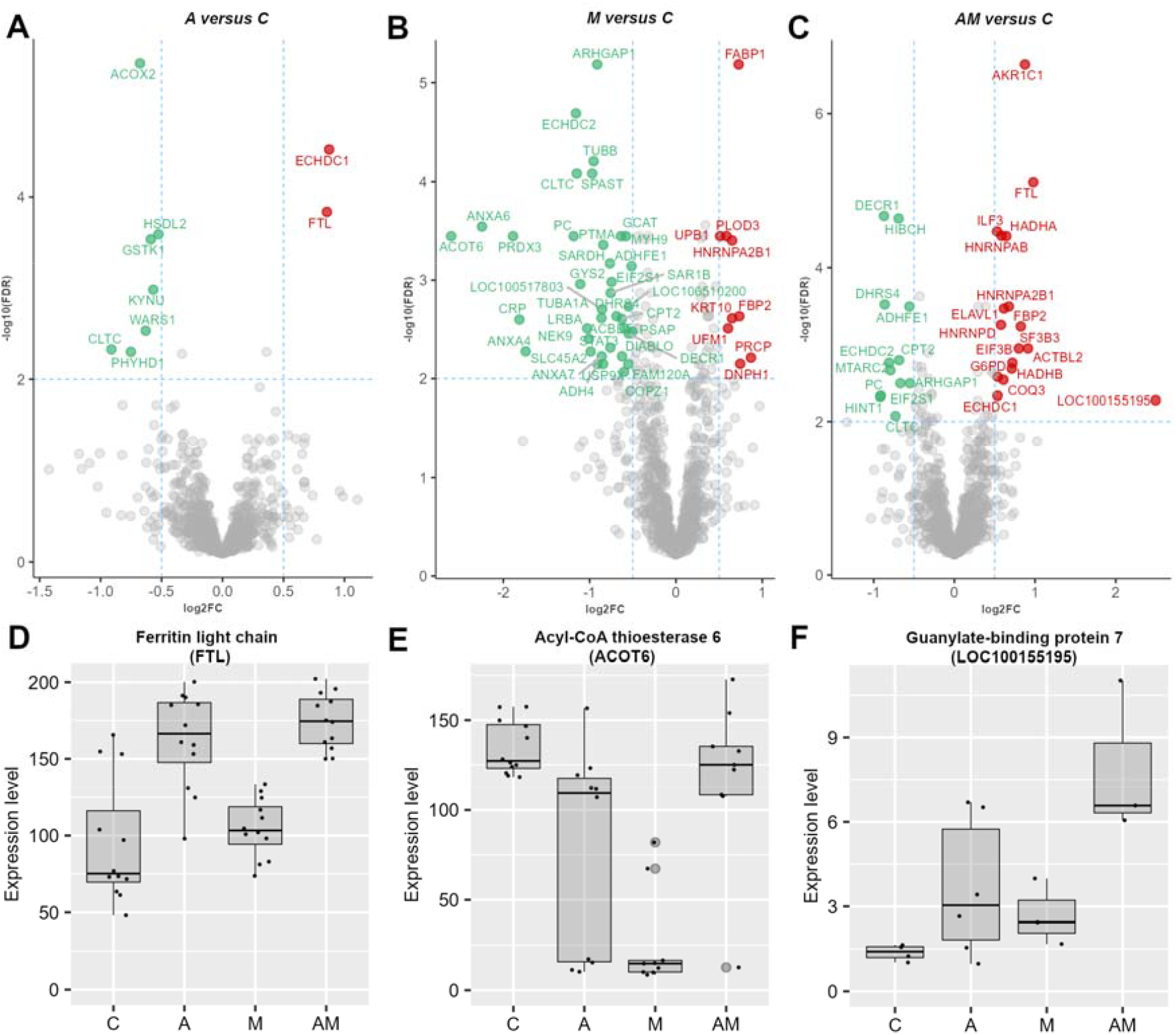
Volcano plots for liver cytoplasmatic fraction, showing log2FC from A versus C (**A**), M versus C (**B**), and AM versus C (**C**), at FDR adjusted *p* value < 0.01, and log2 threshold (-0.5, 0.5). Bar plots represent protein expression level of ferritin light chain (**D**), acyl-CoA thioesterase 6 (**E**) and guanylate-binding protein 7 (**F**) in liver cytoplasmatic samples from control and experimental (A, M and AM) groups. C group represent the control group of weaned piglets (C) fed with a basal diet. The A group were fed with the basal diet plus a mixture of two antioxidant byproducts (grapeseed and sea buckthorn meal. The M group were fed with the basal diet artificially contaminated with two mycotoxins (AFB1 and OTA). The AM group represent the weaned piglets fed with the basal diet containing the mixture (1:1) of antioxidant byproducts and the two mycotoxins.

The log-ratio data from the comparison of the group fed with mycotoxins (M) contaminated diet artificially with AFB1 and the control (C) group (**Figure 1**B) revealed that acyl-CoA thioesterase 6 (ACOT6) (**Figure 1**F) showed the most substantial decrease in expression (log2FC = -2.60), followed by annexin A6 (ANXA6) with a similarly decrease (log2FC = -2.25). However, peroxiredoxin 3 (PRDX3) showed a slightly lower expression (log2FC = -1.89). On the other hand, prolylcarboxypeptidase (PRCP) exhibited the highest increase in expression (log2FC = 0.87), followed by 2’-deoxynucleoside 5’-phosphate N-hydrolase 1 (DNPH1) with a notable increase (log2FC = 0.74), and by fatty acid binding protein 1 (FABP1) with the lowest FDR from the up-regulated proteins (log2FC = 0.72).

When comparing the concomitant administration of antioxidants and mycotoxins (AM) group to the control group (C), we noticed significant changes in expression level of 163 proteins, with 33 having an *p* value < 0.01 (**Figure 1**C). Among these, histidine triad nucleotide binding protein 1 (HINT1) and pyruvate carboxylase (PC) exhibited the most substantial expression reduction (log2FC = -0.92, and log2FC = -0.91, respectively). On the other hand, guanylate-binding protein 7 (LOC100155195) (**Figure 1**F) showed the highest overexpression (log2FC = 2.49). There were also notable declines in the expression of enoyl-CoA hydratase domain containing 2 (ECHDC2), dehydrogenase/reductase (SDR family) member 4 (DHRS4), and 2,4-dienoyl-CoA reductase 1 (DECR1), suggesting potential perturbations in cytoplasmatic metabolic pathways. Furthermore, the expression of ferritin light chain (FTL) aldo-keto reductase family 1, member C1 (AKR1C1) and interleukin enhancer binding factor 3 (ILF3) showed significant increases, suggesting potential compensatory responses or adaptations to the concomitant administration of antioxidants and mycotoxins.

Enrichment analysis of the liver proteins, cytoplasmatic fraction (**Table 1, Figures S2** and **S3**) for A versus C, M versus C, and AM versus C group comparations, supports the findings observed in the volcano plots. In the A versus C comparison, propanoate metabolism exhibited the highest fold enrichment (27.68), with up-regulation of ethylmalonyl-CoA decarboxylase 1 (ECHDC1) and down-regulation of acyl-CoA oxidase 1 (ACOX1) and aldehyde dehydrogenase 6 family member A1 (ALDH6A1). Also, a down-regulation of proteins involved in peroxisome pathway, tryptophan metabolism, lysosome pathway, and biosynthesis of unsaturated fatty acids was observed. Similarly, in the M versus C comparison, valine, leucine, and isoleucine degradation pathway displayed significant fold enrichments, including up-regulation of 3-hydroxy-3-methylglutaryl-CoA synthase 1 (HMGCS1) and down-regulation of acyl-CoA dehydrogenase short chain (ACADS), acyl-CoA dehydrogenase short/branched chain (ACADSB), 3-hydroxyisobutyrate dehydrogenase (HIBADH), aldehyde dehydrogenase 6 family member A1 (ALDH6A1), aldehyde dehydrogenase 7 family member A1 (ALDH7A1), methyl crotonyl-CoA carboxylase subunit 1 (MCCC1), and 3-hydroxy-3-methylglutaryl-CoA lyase (HMGCL). Moreover, propanoate metabolism, beta-alanine metabolism, butanoate metabolism, phenylalanine metabolism, TCA cycle, glyoxylate and dicarboxylate metabolism, and purine metabolism showed higher fold enrichments in M versus C comparison. Additionally, in the AM versus C comparison, fatty acid degradation, pantothenate and CoA biosynthesis, and beta-alanine metabolism, highlighted up-regulation of another member from aldehyde dehydrogenase protein family, respectively aldehyde dehydrogenase 3 family member A2 (ALDH3A2). In addition to the other 2 comparisons (A versus C and M versus C), in the AM versus C comparison, regulation of actin cytoskeleton, adherent’s junction, focal adhesion and PPAR signalling pathways, showed higher fold enrichment figures.

**Table 1.**
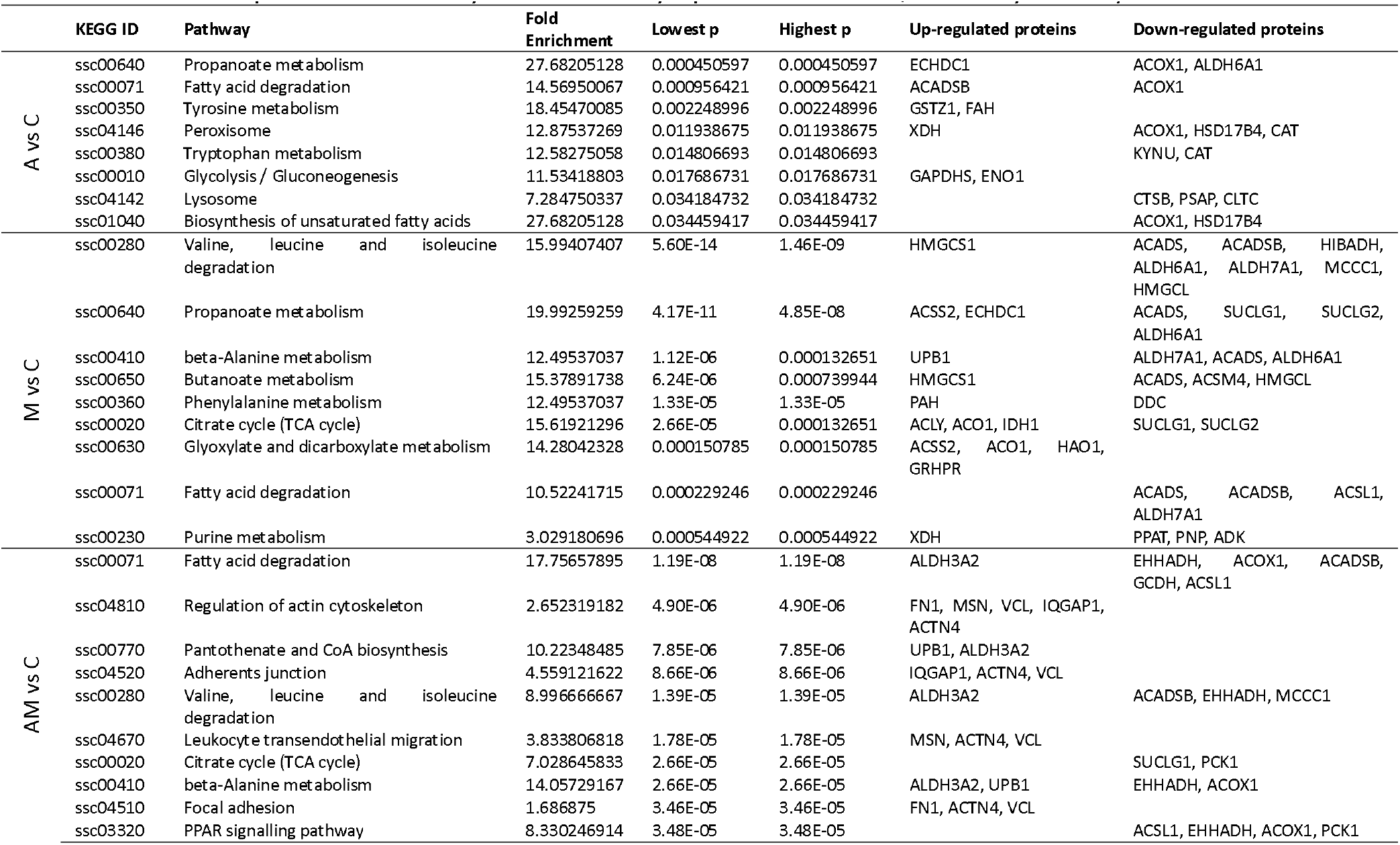
Top 10 KEGG Pathways data in liver cytoplasmatic fraction, sorted by lowest *p* value.

### 3.2. The renal function is impacted by the addition of antioxidants to feed

Metabolites resulting from phase I and II of detoxification are primarily eliminated through urinary excretion, which plays a critical role in the body, with the potential to prevent cell damage and toxicity. Therefore, kidney function and metabolism are directly influenced by the amount and type of mycotoxins present in the diet. In addition to analysing the modifications in the cytoplasmatic fraction of liver, our study also focused on identifying the proteome changes taking place within the kidney, following exposure to antioxidant-enriched diet (A group), mycotoxin-contaminated diet (M group), and their concomitant administration in the diet (AM group).

When kidney cytoplasmatic samples from the A group were compared to those from the control group (C), the log2FC of several proteins changed significantly (**Figures 2**A). Particularly, carnosine dipeptidase 1 (CNDP1), with the lowest *p* value, exhibited a decrease in expression (log2FC = -0.52), as did serine hydroxy-methyltransferase 1 (SHMT1) (log2FC = -0.51). Additionally, carbamoyl-phosphate synthase 1 (CPS1) showed the most decreased expression (log2FC = -2.56). On the other hand, 2’-5’-oligoadenylate synthetase 1 (OAS1) and S100 calcium binding protein A4 (S100A4) demonstrated increased expression with log2FC values of 2.13 and 1.79, respectively. Additionally, several other proteins, including small nuclear ribonucleoprotein D2 polypeptide (SNRPD2), heterogeneous nuclear ribonucleoprotein A/B (HNRNPAB), and ribosomal protein S16 (RPS16), displayed varying degrees of up-regulated expression, suggesting potential roles in the kidney’s response to the antioxidant diet.

**Figure 2.**
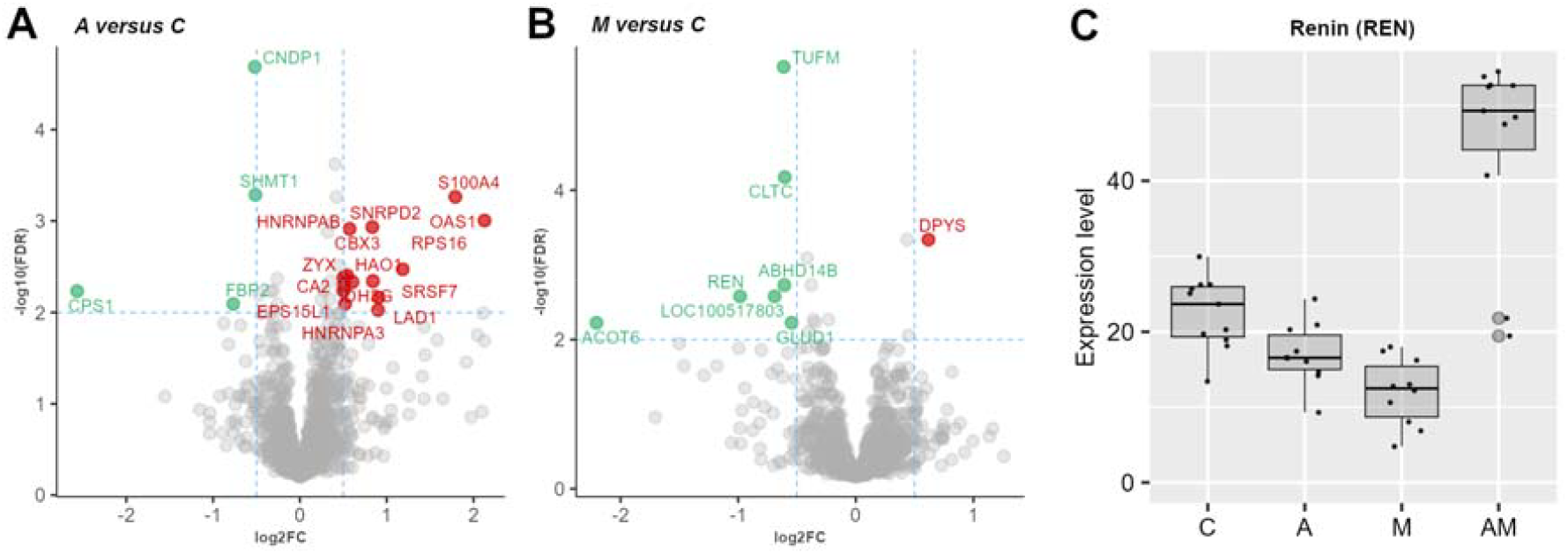
Volcano plots for kidney cytoplasmatic fraction, showing log2FC from A versus C (**A**), and M versus C (**B**), at FDR adjusted *p* value < 0.01, and log2 threshold (-0.5, 0.5). Bar plots represent protein expression level of renin (**C**) in the kidney cytoplasmatic samples from control and experimental (A, M and AM) groups. C group represent the control group of weaned piglets (C) fed with a basal diet. The A group were fed with the basal diet plus a mixture of two antioxidant byproducts (grapeseed and sea buckthorn meal. The M group were fed with the basal diet artificially contaminated with two mycotoxins (AFB1 and OTA). The AM group represent the weaned piglets fed with the basal diet containing the mixture (1:1) of antioxidant byproducts and the two mycotoxins.

The analysis of the log2FC data between the M group and the C group (**Figure 2**B) showed the most substantial decrease in expression for acyl-CoA thioesterase 6 (ACOT6, log2FC = -2.20), followed by renin (REN) with a slightly lower expression (log2FC = -0.98, **Figure 2**C), glycine N-phenyl-acetyltransferase (LOC100517803, log2FC = -0.69), Tu translation elongation factor, mitochondrial (TUFM, log2FC = -0.61), abhydrolase domain containing 14B (ABHD14B, log2FC = --0.61), clathrin heavy chain (CLTC, log2FC = -0.60), and glutamate dehydrogenase 1 (GLUD1, log2FC = -0.55). On the other hand, dihydropyrimidinase (DYPS) is the only up-regulated protein (log2FC = 0.61).

As in the case of liver samples, we observed significant differences between the AM group and the control group (C), for a larger number of proteins, specifically 77 proteins (**Figures 3**A and **S4**). Among these, spindle apparatus coiled-coil protein 1 (SPDL1, log2FC = -1.29), fatty acid binding protein 1 (FABP1, log2Fc = -1.26), ribosomal protein L17 (RPL17, log2FC = -1.07), cingulin like 1 (CGNL1, log2FC = -0.98, **Figure 3**B), and clathrin interactor 1 (CLINT1, log2FC = -0.91, **Figure 3**C) showed the most down-regulation effect. On the other hand, 2’-5’-oligoadenylate synthetase 1 (OAS1, **Figure 3**D) showed the highest upregulation (log2FC = 1.51). There were also up-regulation in ISG15 ubiquitin like modifier (ISG15, log2FC = 1.48), thymosin beta 10 (TMSB10, log2FC = 1.10), renin (REN, log2FC = 0.95, **Figure 2**C), fructose-bisphosphatase 2 (FBP2, log2FC = 0.90), and glyceraldehyde-3-phosphate dehydrogenase (GAPDH, log2FC = 0.76, **Figure 3**E). All log-ratios of identified proteins in the kidney cytoplasmatic samples are provided in **Table S2**).

**Figure 3.**
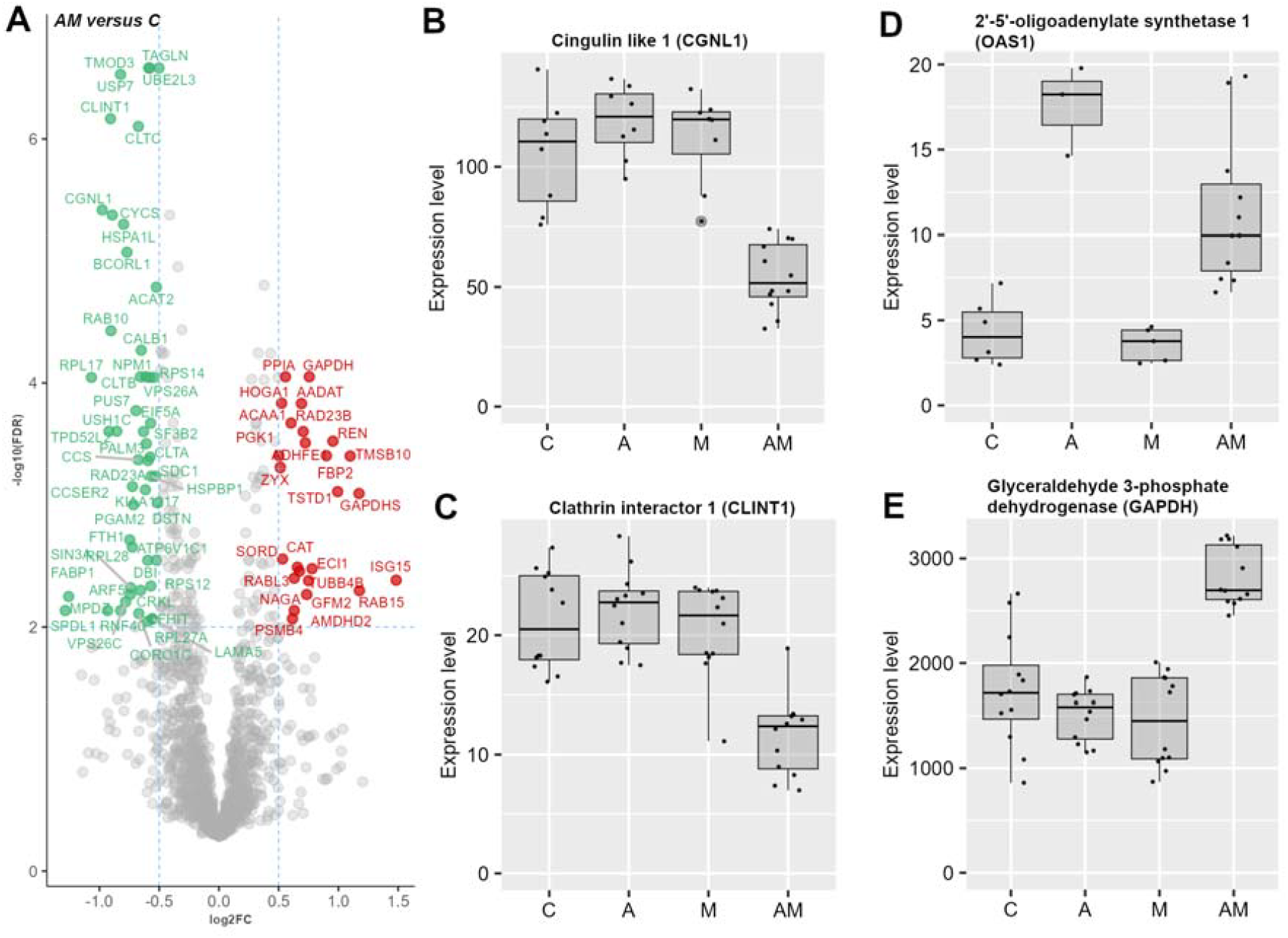
Proteomic changes in the kidney cytoplasmatic fraction induced by concomitant administration of antioxidants and mycotoxins in the piglet’s diet. Volcano plots showing log2FC from AM versus C (**A**), at FDR adjusted *p* value < 0.01, and log2 threshold (-0.5, 0.5). Bar plots represent protein expression level of cingulin like 1 (**B**), clathrin interactor 1 (**C**), 2’-5’-oligoadenylate synthetase 1 (**D**), and glyceraldehyde-3-phosphate dehydrogenase (**E**) in the kidney cytoplasmatic samples from control and experimental (A, M and AM) groups. C group represent the control group of weaned piglets (C) fed with a basal diet. The A group was fed with the basal diet plus a mixture of two antioxidant byproducts (grapeseed and sea buckthorn meal. The M group was fed with the basal diet artificially contaminated with two mycotoxins (AFB1 and OTA). The AM group represents the weaned piglets fed with the basal diet containing the mixture (1:1) of antioxidant byproducts and the two mycotoxins.

Enrichment analysis (**Table 2, Figures S5** and **S6**) suggests that exposure to antioxidant-, mycotoxin-contaminated, and combining antioxidant byproducts and AFB1 and OTA mycotoxins diet, induces significant changes in the piglet’s kidney cytoplasmatic proteome, specifically in the expression of proteins involved in amino acid synthesis and metabolism, and catabolism, like fatty acid degradation, glycolysis, valine, leucine and isoleucine degradation and TCA cycle. These results offer details about the molecular mechanisms that induce alterations in renal amino acid metabolism when combined exposure of antioxidants and mycotoxins took place, compared to exposure to either antioxidants or mycotoxins alone. Therefore, in the A versus C comparison, amino acid metabolism, including beta-alanine, arginine and histidine pathways showed higher fold enrichments, through up-regulation of aldehyde dehydrogenase 1 family member B1 (ALDH1B1), and aminoacylase 1 (ACY1), and down-regulation of CNDP1 (**Figure 2**A), and beta-ureidopropionase 1 (UPB1). In the M versus C comparison, beta alanine metabolism exhibited the highest fold enrichment (27.21), with up-regulation of CNDP2, and UPB1 and down-regulation of ALDH1B1, enoyl-CoA hydratase and 3-hydroxyacyl CoA dehydrogenase (EHHADH), ALDH6A1. Also, a down-regulation of proteins involved in valine, leucine and isoleucine degradation, TCA cycle, pentose phosphate pathway, and fatty acid degradation was observed. Catalase (CAT) expression was observed to be up-regulated in several pathways, respectively in peroxisome, glycolysis and glyoxylate and dicarboxylate metabolism. On the other hand, in the AM versus C comparison, valine, leucine, and isoleucine degradation pathway displayed up-regulation of ACADSB, EHHADH, acetyl-CoA acyltransferase 1 (ACAA1, **Figure 3**A), propionyl-CoA carboxylase subunit alpha (PCCA), ALDH6A1, and down-regulation of acetyl-CoA acetyltransferase 2 (ACAT2). Moreover, proteasome, TCA cycle, fatty acid degradation, fructose and mannose metabolism, protein processing in endoplasmic reticulum pathway, cysteine and methionine metabolism, and glycolysis pathways, showed both up- and down-regulated proteins in AM versus C comparison.

**Table 2.**
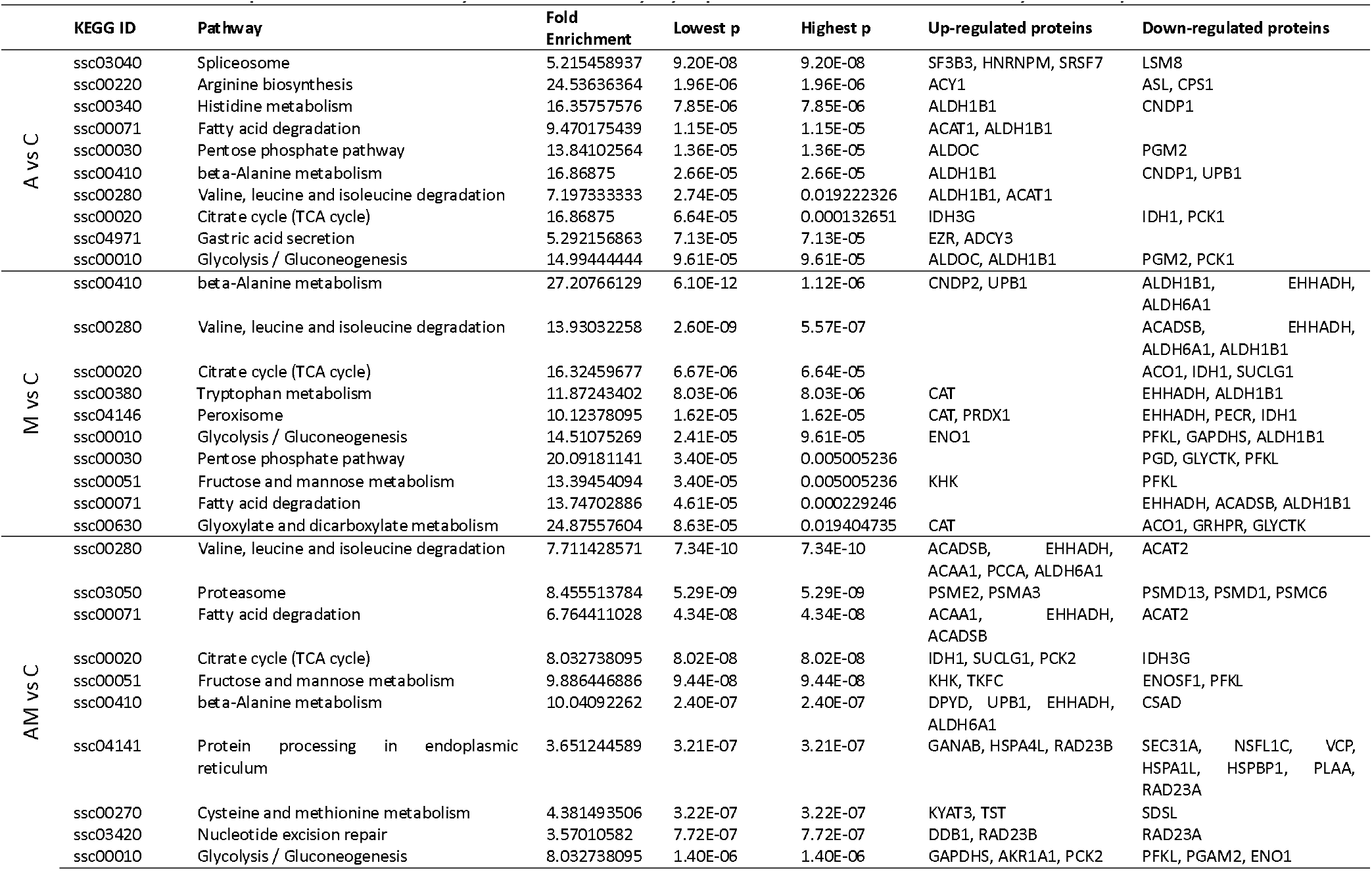
Top 10 KEGG Pathways data in kidney cytoplasmatic fraction, sorted by lowest *p*.

### 3.3. Impact of mycotoxins

To investigate the impact of mycotoxins by AFB1 and OTA exposure in conjunction with antioxidants supplementation compared to antioxidant intake alone (AM versus A), we conducted a comparative differential expression analysis of proteins that meets the criteria of *p* value < 0.01 and FC outside (-1, 1) interval in liver and kidney cytoplasmatic fractions. In the liver cytoplasmatic fraction, only one spliceosome member, splicing factor 3b subunit 3 (**Figure 4A**), showed significant up-regulation (log2FC = 1.03). Applying a less stringent fold change interval of (-0.5, 5) to the liver data, eight proteins were identified (**Figure S7**). These proteins include kynureninase (KYNU), hydroxyacyl-CoA dehydrogenase trifunctional multienzyme complex subunit alpha (HADHA), SF3B3, aldo-keto reductase family 1, member C1 (AKR1C1), tryptophanyl-tRNA synthetase 1 (WARS1), and brain abundant membrane-attached signal protein 1 (BASP1) in the first cluster, and superoxide dismutase 2 (SOD2) and 3-hydroxyisobutyryl-CoA hydrolase (HIBCH) in the second cluster.

**Figure 4.**
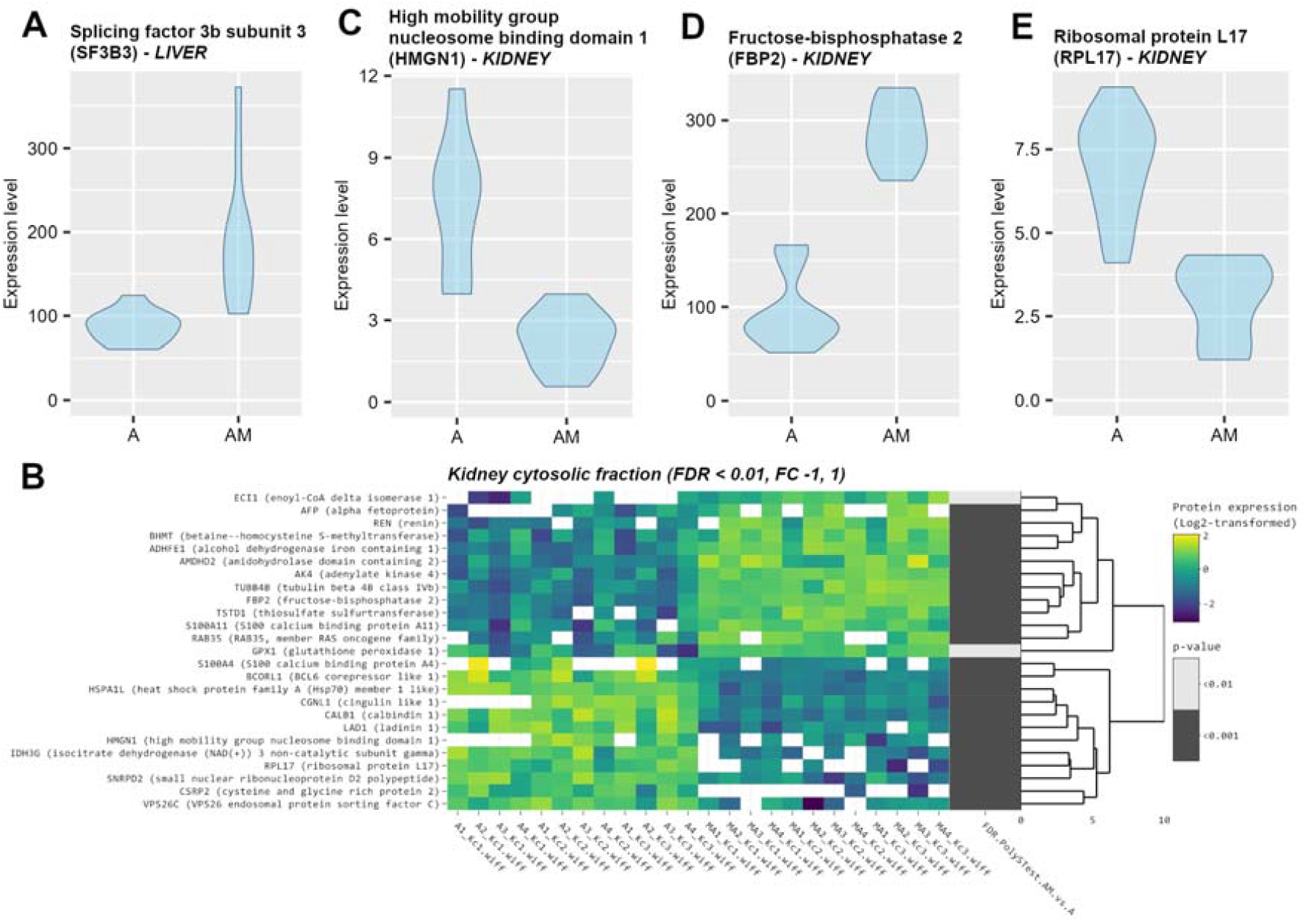
Impact of mycotoxins in the piglet’s diet. Violin plots represent protein expression level of splicing factor 3b subunit 3 (**A**) from liver cytoplasmatic samples from A and AM groups. Clustered heatmap of the 25 differentially expressed proteins, filtered with log2FC threshold set to exclude interval (-1, 1) and FDR adjusted *p* value < 0.01. Yellow colour represents upregulation, while blue represents downregulation in A group compared to AM group. From left to right, expression values (log2 transformed) for replicates (4 biological × 3 technical) are shown for the A group and for the AM group, followed by significance values of the comparison to A group (**B**). Violin plots represent protein expression level of high mobility group nucleosome binding domain 1 (**C**), fructose-bisphosphatase 2 (**D**), and ribosomal protein L17 (**E**) in kidney cytoplasmatic samples from A and AM groups. The A group was fed with the basal diet plus a mixture of two antioxidant byproducts (grapeseed and sea buckthorn meal. The AM group represent the weaned piglets fed with the basal diet containing the mixture (1:1) of antioxidant byproducts and the two mycotoxins.

On the other hand, the impact of mycotoxins on the cytoplasmatic fraction in the kidneys is much more significant, the degree of protein expression level alteration being much higher, with 25 proteins with log 2-fold change outside interval (-1, 1) (**Figure 4B**), also grouped in two clusters. The first cluster include up-regulated proteins that are related to metabolic homeostasis, including enzymes involved in gluconeogenesis, fructose-bisphosphatase 2 (FBP2, **Figure 4D**), cytoskeletal organization through tubulin beta 4B class IVb (TUBB4B), maintenance of energy, by adenylate kinase 4 (AK4), amidohydrolase domain containing 2 (AMDHD2), betaine-homocysteine S-methyltransferase (BHMT), and renal function through renin (REN) and alpha fetoprotein (AFB). The second cluster contains down-regulated proteins primary involved in nucleosome binding and ribosomal function, high mobility group nucleosome binding domain 1 (HMGN1, **Figure 4C**) and ribosomal protein L17 (RPL17, **Figure 4E**), respectively, but also chaperones like heat shock protein family A (Hsp70) member 1 like (HSPA1L), proteins involved in cytoskeletal organization such as calbindin 1 (CALB1), calcium homeostasis (calbindin 1 – CALB1), TCA cycle (isocitrate dehydrogenase (NAD(+)) 3 non-catalytic subunit gamma – IDH3G), endosomal trafficking (VPS26 endosomal protein sorting factor C – VPS26C), nucleic acid binding proteins like BCL6 corepressor like 1 (BCORL1), and others like cingulin like 1 (CGNL1), ladinin 1 (LAD1), small nuclear ribonucleoprotein D2 polypeptide (SNRPD2), cysteine and glycine rich protein 2 (CSRP2), and S100 calcium binding protein A4 (S100A4).

### 3.4. Impact of antioxidants

The influence of addition of antioxidants in the diet contaminated with mycotoxins (AM versus M), was observed only in the kidney cytoplasmatic samples, at log2 fold change outside the range (-1, 1) (**Figure 5**A), while in the liver cytoplasmatic fraction, the protein expression changes were not so high (**Figure S**8). Thus, 20 proteins with log2FC < -1 and >1 were identified, grouped in 2 clusters with 14 up-regulated proteins in the first cluster and 6 down-regulated proteins in the 2nd cluster. The heatmap (**Figure 5**B) shows that some of the up-regulated proteins in the first cluster, namely ADHFE1 (log2FC = 1.01), AK4 (log2FC = 1.16), BHMT (log2FC = 1.24), ECI1 (log2FC = 1.03), RAB35 (log2FC = 1.05), and TSTD1 (log2FC = 1.05), presented similar log2FC values to those observed as the effect of mycotoxins (**Figure 4**B). The same pattern is observed for the down-regulated proteins from the second cluster, more precisely for: CGNL1 (log2FC = -1.07), HMGN1 (log2FC = -1.79) and RPL17 (log2FC = -1.23).

**Figure 5.**
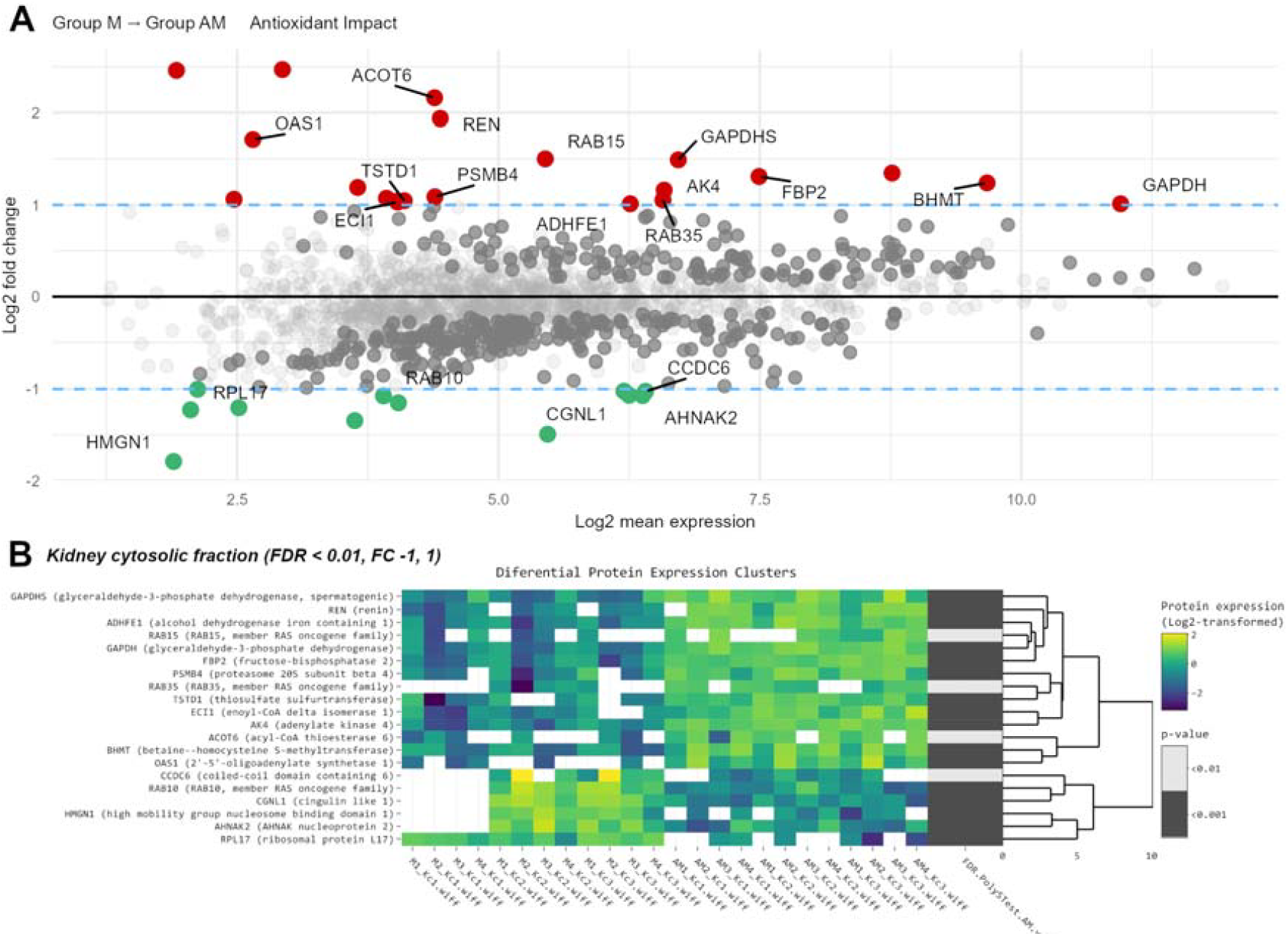
Impact of antioxidants in the piglet’s diet. MA plot showing log2FC versus log2 mean expression from AM versus M comparison. Red colour represents up-regulated proteins, and green colour represent down-regulated proteins, log2FC outside interval (-1, 1). Dark grey colour represents significant data with FDR adjusted *p* value < 0.01 (**A**). Clustered heatmap of the 20 differentially expressed proteins, filtered with log2FC outside (-1, 1). Yellow colour represents upregulation, while blue represents downregulation in M group compared to AM group. From left to right, expression values (log2 transformed) for replicates (4 biological × 3 technical) are shown for the M group and for the AM group, followed by significance values of the comparison to M group (**B**). The M group was fed with the basal diet artificially contaminated with two mycotoxins (AFB1 and OTA). The AM group represents the weaned piglets fed with the basal diet containing the mixture (1:1) of antioxidant byproducts and the two mycotoxins.

FBP2 showed a lower expression level (log2FC = 1.31) while REN was up-regulated with 41.60% (log2FC = 1.94). Also, in the first cluster, some up-regulated proteins were observed, whose expression seems specific to the effect given by the presence of antioxidants, such as proteins involved in lipid metabolism, like acyl-CoA thioesterase 6 (ACOT6, log2FC = 2.16), involved in energetic metabolism, like glyceraldehyde-3-phosphate dehydrogenase (GAPDH, log2FC = 1.01) and GAPDHS (GAPDH, spermatogenic, log2FC = 1.49), involved in protein turnover, like proteasome 20S subunit beta 4 (PSMB4, log2FC = 1.09), and implicated in the intracellular vesicle trafficking, like RAB15, member RAS oncogene family (log2FC = 1.50). Moreover, down-regulated proteins, in the second cluster, like AHNAK nucleoprotein 2 (AHNAK2, log2FC = -1.07), coiled-coil domain containing 6 (CCDC6, log2FC = -1.02) and RAB10, member RAS oncogene family (RAB10, log2FC = -1.07) were observed.

## 4. Discussion

The balance between detoxification and toxic action depends on the dose and the possibility of saturation of metabolic pathways, individual genetic variation, or species. Tissue differences in the isoenzyme pattern are influenced by age, diet, health status and sex. In this study we aimed to shed light on the potential benefits of antioxidant supplementation in mitigating mycotoxin toxicity on liver and kidney function, focusing on changes in the cytoplasmatic proteome.

In the liver cytoplasmatic fraction, the addition of antioxidants in diet (A versus C) and piglets’ exposure to mycotoxin-contaminated feed (M versus C), show impact on proteins expression for acyl-CoA oxidase 2 (ACOX2), phytanoyl-CoA dioxygenase domain containing 1 (PHYHD1), acyl-CoA thioesterase 6 (ACOT6), and annexin A6 (ANXA6) (**Figure 1**B), which are involved in fatty acid metabolism, peroxisome pathway, and exosome (**Table 1, Figure S9**). These findings align with mycotoxins metabolism, through the xenobiotic biotransformation pathway, involving their conversion and intracellular translocation, allowing the transformation of toxic substances into compounds suitable for excretion (Esteves et al. 2021). Therefore, the detoxification process occurs in three phases: modification, or phase I, characterised by addition of functional groups; conjugation or phase II, involving the conjugation of functional groups; and excretion or phase III. Common phase I reactions include oxidation, reduction, hydrolysis, hydration, and dehalogenation. During this phase, mycotoxins are initially transformed into more water-soluble chemical form that can be metabolized by the enzymes involved in phase II (Croom 2012; Gerdemann et al. 2023). As we previously observed by proteomic analysis of the microsomal fraction from the liver (Popescu et al. 2023), cytochromes P450 isoforms have a central role in catalysing phase I reactions.

The co-administration of antioxidants and mycotoxins (AM versus C) induced significant changes in the expression of several proteins in liver cytoplasmatic fraction. Histidine triad nucleotide binding protein 1 (HINT1) is involved in DNA repair mechanisms (Wei et al. 2018). Therefore, its down-regulation may indicate potential disruptions in this process. Pyruvate carboxylase (PC) plays a central role in hepatic gluconeogenesis and lipogenesis by catalysing the carboxylation of pyruvate to oxaloacetate, a key intermediate in glucose and lipid metabolism (Jitrapakdee et al. 2008). Additionally, PC contributes to restoration of the citric acid cycle intermediates and maintains metabolic flux in the liver (Kiesel et al. 2021). The reduction in PC protein expression (**Figure S10**) suggests a potential impairment in gluconeogenesis and lipogenesis, which are essential for energy production and lipid metabolism in the liver. Guanylate-binding protein 7 (LOC100155195) belongs to the guanylate-binding protein (GBP) family, and may indicate activation of immune responses to environmental stimuli (Feng et al. 2021), suggesting that the liver is undergoing dynamic changes to counteract potential side effects and to maintain tissue homeostasis. Enrichment analysis further supports these findings, showing changes in pathways related to propanoate metabolism (**Figure S11**), fatty acid degradation (**Figure S12**), and others (**Figures S13 – S18**), indicating the complex regulatory mechanisms that support liver function in response to dietary factors.

Phase II reactions, such as glucuronidation, sulfation, and conjugation with glutathione (GSH) occur primally in the cytoplasm and are the most important in xenobiotic toxicity. The cofactors utilised in these conjunction reactions are uridine 5⍰-diphosphoglucuronic acid (UDPGA), 3⍰-Phosphoadenosine-5⍰-phosphosulfate (PAPS) and GSH. UDP-glucuronosyltransferases (microsomal) and sulfotransferases (cytoplasmatic) catalyse glucuronidation and sulfation reactions, respectively. These enzymes have tissue and species-specific isoforms. For example, in pigs’ liver, specific markers for UDP-glucuronosyltransferases and sulfotransferases are UGT1A1, UGT1A3, UGT1A6, UGT1A10, UGT2B18-like, UGT2B31, UGT2B31-like, SLUT1A1 and SLUT2A1. The resulting glucuronide and sulphate conjugates increase the water solubility of the parent compounds and facilitate their excretion through urine and bile. However, there are cases where phase II detoxification reactions can lead to increased toxicity, as seen in bilirubin glucuronidation by UGT1A1. For instance, fumonisin B1 competes with bilirubin for UGT1A1, leading to hyperbilirubinemia due to enzyme activity inhibition by substrate competition. According to our results, the most affected protein expression levels associated with glucuronidation and sulfation included acyl-CoA thioesterase 6 (ACOT6), aldo-keto reductase family 1 member C1 (AKR1C1), renin (REN), and fatty acid binding protein 1 (FABP1). Other phase II reactions include acetylation catalysed by N-acetyltransferases, methylation of nitrogen (N), oxygen (O), sulphur (S), and arsenic (As) mediated by methyltransferases, and conjugation with amino acids (Miyagi and Collier 2011; Yang et al. 2017; Mathur et al. 2001; Jančová et al. 2010; Lehman-McKeeman et al. 2018).

Moreover, phase II reactions provide cellular protection against oxidative stress through the action of antioxidant enzymes: CAT, SOD, and GST (**Figure S9**). However, under conditions of increased toxicity, the products of phase II may induce changes in protein expression, reactive oxygen species (ROS) production, and oxidative stress in the affected cells (Jarolim et al. 2016; Wen et al. 2016; Antonissen et al. 2017; Ngo and Duennwald 2022). Also, our differential analysis highlights specific proteins involved in antioxidant defence mechanisms and ROS production (**Figure S9**), such as catalase (CAT), renin (REN), hydroxysteroid dehydrogenase like 2 (HSDL2), glutathione S-transferase kappa 1 (GSTK1), and fatty acid binding protein 1 (FABP1).

On the other hand, in the kidney cytoplasmatic fraction, our results revealed a significant decrease in proteins expression associated with amino acid metabolism in response to the antioxidant-enriched diet (**Table 2,** **Figure 2**A). Specifically, down-regulation of carnosine dipeptidase 1 (CNDP1), involved in arginine, proline (**Figure S19**), histidine (**Figure S20**), and beta-alanine metabolism (**Figure S21**); serine hydroxy-methyltransferase 1 (SHMT1), an essential enzyme in carbon pool by folate (**Figure S22**), and biosynthesis of amino-acids; and carbamoyl-phosphate synthase 1 (CPS1), a key enzyme nitrogen metabolism and urea cycle (**Figure S23**), indicate alterations in these metabolic pathways. In contrast, we observed increased expression of 2’-5’-oligoadenylate synthetase 1 (OAS1), a protein involved in the immune response, and S100 calcium binding protein A4 (S100A4), often associated with inflammation and cell migration, that might indicate a response to the antioxidant diet.

Additionally, comparison between the mycotoxin-contaminated diet group (M group) and the control group (C) (**Table 2,** **Figure 2**B) showed a significant down-regulation of acyl-CoA thioesterase 6 (ACOT6), renin (REN), and other proteins, along with up-regulation for dihydropyrimidinase (DYPS). ACOT6 catalyses the hydrolysis of acyl-CoA thioesters to free fatty acids and coenzyme A, regulating intracellular levels of fatty acids (Brocker et al. 2010). In the kidney, ACOT6 is involved in lipid biosynthesis, maintaining lipid homeostasis. Renin, generated and secreted by kidney juxtaglomerular cells, is a key enzyme in the renin-angiotensin-aldosterone system (RAAS), and it regulates blood pressure and fluid-electrolyte balance by catalysing the conversion of angiotensinogen to angiotensin I, which produces vasoconstrictor angiotensin II. Aldosterone, released by angiotensin II, promotes sodium and water retention, that influence blood pressure and kidney function (Broeker et al. 2022). While the role of DYPS in the kidney is not fully understood, it is involved in nucleic acid synthesis (**Figure S24**) and beta-alanine metabolism (**Figure S21**). These changes may indicate an adaptive response of the kidney to mitigate the effects of mycotoxin-induced stress, or direct toxicity to renal cells, highlighting the intricate interplay between environmental exposures and renal function.

When the concomitant administration of antioxidants and mycotoxins (AM group) was evaluated (**Figure 3**), significant alterations in protein expression were observed. This included down-regulation of spindle apparatus coiled-coil protein 1 (SPDL1), and fatty acid binding protein 1 (FABP1), along with the up-regulation of OAS1 and REN, suggesting complex interactions between antioxidants and mycotoxins. In the kidney cytoplasmatic fraction, SPDL1 contributes to the maintenance of genomic stability during cell division (Song et al. 2022), ensuring proper development and function of renal cells. FABP1, primarily expressed in the liver but also found in renal proximal tubular cells, is involved in the PPAR signalling pathway (**Figure S25**), through uptake, transport, and utilization of fatty acids, which are essential for energy production and membrane synthesis in renal cells (Wang et al. 2015). The down-regulation of SPDL1 and FABP1 expression might indicate disruptions in the cell division processes and lipid metabolism, respectively, while the up-regulation of REN suggests potential involvement in the urea cycle. Enrichment analysis further highlighted changes in pathways associated with amino acid metabolism, fatty acid degradation (**Figure S26**), and glycolysis (**Figure S27**), indicating a multifaceted impact of dietary factors on kidney function and metabolism.

The body eliminates unwanted substances by excretion, which plays an important role in preventing cellular damage and toxicity. Metabolites resulting from phase I and II of detoxification are mainly excreted via urine and bile, while gases are eliminated by exhalation. Also, xenobiotic metabolites can be excreted through sweat, saliva, tears, and breast milk (Lehman-McKeeman et al. 2018; Badrigilan et al. 2020). Although the kidneys constitute around 4% of body mass, they receive almost 25% of cardiac output, representing the major route in eliminating compounds and electrolytes via urinary excretion. Glomerular filtration is influenced by the pressure differences between the afferent and efferent arterioles, capillaries pore size, and the degree of xenobiotic binding to plasma proteins. Therefore, glomerular filtration allows passage of molecules below 60 kDa. Which depends on the chemical nature of the compounds undergoing excretion and the number of available nephrons. Once filtered, a compound may be excreted or reabsorbed, mainly in the proximal tubules by passive diffusion, depended by metabolite solubility and ionization. Additionally, xenobiotics can be excreted by an active mechanism mediated by transporters (Brater 2002; Ruggiero et al. 2010; Chapman et al. 2020). According to our results, proteins such as renin, acyl-CoA thioesterase 6, and fatty acid binding protein 1 (FABP1) from the kidney cytoplasmatic fraction may play a role in the transport of xenobiotics.

The impact of mycotoxin exposure (AFB1 and OTA) along with antioxidant supplementation (AM versus A) revealed distinct changes in protein expression levels within liver and kidney cytoplasmatic fractions. In liver, only the splicing factor 3b subunit 3 (SF3B3) (**Figure 4**A), a complex responsible for pre-mRNA splicing, showed significant upregulation, which may affect mRNA splicing patterns, and potentially affecting the expression of genes involved in stress response. Following a less stringent fold change interval, kynureninase (KYNU), hydroxyacyl-CoA dehydrogenase trifunctional multienzyme complex subunit alpha (HADHA), superoxide dismutase 2 (SOD2), and 3-hydroxyisobutyryl-CoA hydrolase (HIBCH), showed significant metabolic adaptations (**Figure S7**). KYNU catalyses the conversion of 3-hydroxy-L-kynurenine to 3-hydroxyanthranilate, a key step in the degradation of tryptophan (**Figure S14**). Additionally, tryptophanyl-tRNA synthetase 1 (WARS1) catalyses the attachment of tryptophan to its corresponding tRNA, ensuring accurate incorporation of tryptophan into nascent polypeptide chains during translation (Ahn et al. 2021). Thus, WARS1 down-regulation may impact translation as response to changes in tryptophan availability. HADHA catalyses the third step of β-oxidation, converting 3-hydroxyacyl-CoA to 3-ketoacyl-CoA. HADHA expression in liver cells is restored when mycotoxins and antioxidants are co-administrated (**Figures S12, S16**, and **S17**).

Renal function was severely impacted by the presence of mycotoxins, with significant protein expression changes. Up-regulation of FBP2 shows an enhanced capacity for gluconeogenesis, possibly to maintain glucose homeostasis in response to mycotoxins and antioxidant exposure (**Figure 4**D). Also, up-regulation of REN (**Figure 2**C) may suggest activation of RAAS and a role in metabolic homeostasis. Down-regulation of HMGN1 (**Figure 4**C), a chromatin-associated protein that modulates chromatin structure and gene expression by binding to nucleosomes (Zhang et al. 2016), could suggest an impact in DNA replication, repair, and transcription. This observation is supported by the down-regulation of RPL17 (**Figure 4**E), a component of the ribosome (**Figure S28**), which suggest an impairment in the protein synthesis machinery. Furthermore, proteins involved in cytoskeletal organization, chaperones, calcium homeostasis, and nucleic acid binding were affected (**Figure 4**B), demonstrating the complex cellular response to combined dietary exposures in renal function.

The impact of antioxidants in the diet contaminated with mycotoxins (AM versus M) was mainly observed in kidney cytoplasmatic samples. Specifically, several proteins showed similar expression changes to those induced by mycotoxins exposure alone, suggesting overlapping effects. Therefore, ADHFE1 and ECI1, involved in fatty acid degradation (**Figure S26**); AK4, responsible to balance purine metabolism (**Figure S24**); BHMT, an enzyme involved in methylation reactions of phase II (Feng et al. 2011) and in cysteine and methionine metabolism (**Figure S29**); RAB35, involved in intracellular vesicle trafficking (Martinez-Arroyo et al. 2021), and TSTD1, with role in the sulphide oxidation pathway (Libiad et al. 2018); may reflect adaptations to maintain energy homeostasis, and alterations in response to oxidative stress. However, FBP2 and REN showed unique expression profiles, indicating a potential specific effect of antioxidants (**Figure 5A**). Furthermore, the up-regulated proteins (**Figure 5**B) were linked to lipid metabolism (ACOT6), energy metabolism (GAPDH, **Figure S27**), protein turnover (proteasome 20S subunit beta 4 – PSMB4, **Figure S30**), and RAB15 with role in membrane traffic from the early endosome to the recycling endosome (Hutagalung and Novick 2011), indicating diverse cellular responses to antioxidant exposure. On the other hand, down-regulated proteins in the second cluster (**Figure 5**B), including AHNAK nucleoprotein 2 (AHNAK2), coiled-coil domain containing 6 (CCDC6) and RAB10, member RAS oncogene family (RAB10), highlighted additional pathways affected by antioxidant intervention, respectively immune response (Zardab et al. 2022), DNA damage checkpoints (Cerrato et al. 2018), and vesicular trafficking (Tavana et al. 2018). These down-regulation highlights the antioxidants capacity to counteract mycotoxins effects and prevent tumoral cell progression.

The identified effects of antioxidants, as revealed through the alterations in protein expression profiles within the kidney cytoplasmatic fraction, highlight a direct relationship between dietary components and metabolic balance. Although, some proteins showed unique expression patterns in the presence of the antioxidants alone, potentially indicating specific effects, the overlapping changes with exposure to mycotoxins suggest a synergic interaction within renal cellular processes.

The results of this study highlight the need to understand the effects determined by supplementing the diet with a mixture of antioxidants rather than a single antioxidant. Cellular metabolism acts differently when exposed to mixtures compared to individual treatments, as the synergistic and /or antagonistic effects are not well understood. To propose solutions that could mitigate or counteract the effects of the mycotoxin’s contamination in the animal feed, it is important to understand the big picture regarding mechanisms and interactions.

However, our findings are subject to limitations, mainly by the concentrations of mycotoxins and by-products containing antioxidants and included in the diet. Additionally, the present study only evaluated the effects of AFB1 and OTA mycotoxins. The biotransformation of xenobiotics depends on the compound structure, physicochemical properties, and the available enzymes in the exposed tissue. Therefore, future research is needed to confirm dose-dependent effects and explore the influence of other types of mycotoxins and antioxidants.

## 5. Conclusions

This study started from the hypothesis that antioxidants can mitigate the adverse effects of mycotoxins contamination on weaned piglets. It was observed that the effects of mycotoxins are partially mitigated by the antioxidants enriched diet. Our findings show an expression balance of the proteins involved in the amino acids metabolism and antioxidant defense system, accompanied by an energetic compensation through glycolysis and gluconeogenesis. Additionally, we show that in kidneys, some of the effects are synergistically amplified, such as proteins involved in the fatty acids’ degradation, peroxisome, PPAR signaling, translation, TCA cycle and the excretion pathways. Inclusion of antioxidants in the animal diet can have beneficial effects. Nevertheless, caution is advised, synergistic effects can occur with potentially more serious consequences than the effect of mycotoxins alone.

## Supplementary material

**Table S1**. Pig liver cytoplasmatic fraction – quantitative proteomic data.

**Table S2**. Pig kidney cytoplasmatic fraction – quantitative proteomic data.

**Document S1**. Figures S1-S30.

## Acknowledgements

This study was supported by Independent Research Association, Bucharest - 012416, Romania. We are deeply grateful to Dr. Ionelia Țăranu and Dr. Daniela Marin, from the National Research-Development Institute for Animal Nutrition and Biology (Balotești, Romania), for conducting the animal experiment and providing liver and kidney tissue samples for proteomic analysis.

## Ethics

The study was conducted according to the guidelines of the Declaration of Helsinki, and approved by the Ethical Committee of the National Research-Development Institute for Animal Nutrition and Biology, Balotești, Romania (Ethical Committee no. 118/02.12.2019) and in accordance with the Romanian Law 206/2004 and the EU Council Directive 98/58/EC for handling and protection of animals used for experimental purposes.

## Data accessibility

The mass spectrometry proteomics data have been deposited to the ProteomeXchange Consortium via the PRIDE partner repository with the dataset identifier PXD050835.

## Author Contributions

Conceptualization, R.G.P., A.D. and G.C.M.; Methodology, R.G.P and G.C.M; Investigation, R.G.P. and G.C.M.; Writing – Original Draft, R.G.P.; Writing – Review & Editing, G.C.M. and A.D.; Funding Acquisition, G.C.M.; Resources, G.C.M.; Supervision, G.C.M. and A.D.

