## Supplementary material for "Proteomic Changes in the Cytoplasmatic Fraction of Weaned Piglets Liver and Kidney Under Antioxidants and Mycotoxins Diets": Document_S1.pdf

Figure S1

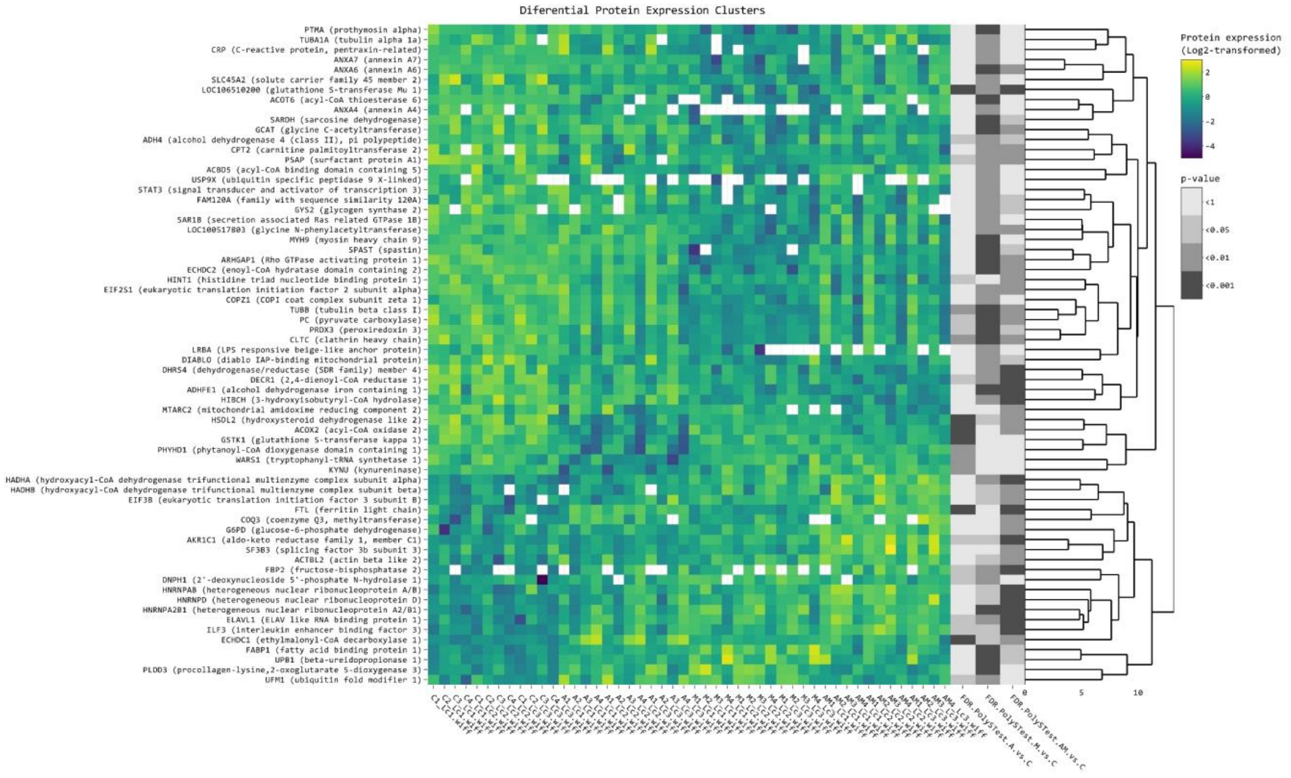


**Figure S1**. Clustered heatmap of the differentially expressed proteins in liver cytoplasmatic fraction, filtered with log2FC threshold set to exclude interval (-0.5, 0.5) and FDR adjusted *p* value < 0.01. Yellow colour represents upregulation, while blue represents downregulation in A, M, and AM group compared to Control (C) group. From left to right, expression values (log2 transformed) for replicates (4 biological × 3 technical) are shown for the A, M, and AM groups, followed by significance values of the A versus C, M versus C, and AM versus C comparisons. The control group (C) was fed with a standard diet for starter piglets. The A group were fed with the basal diet plus a mixture of two antioxidant byproducts (grapeseed and sea buckthorn meal. The M group were fed with the basal diet artificially contaminated with two mycotoxins (AFB1 and OTA). The AM group represent the weaned piglets fed with the basal diet containing the mixture (1:1) of antioxidant byproducts and the two mycotoxins.

Figure S2

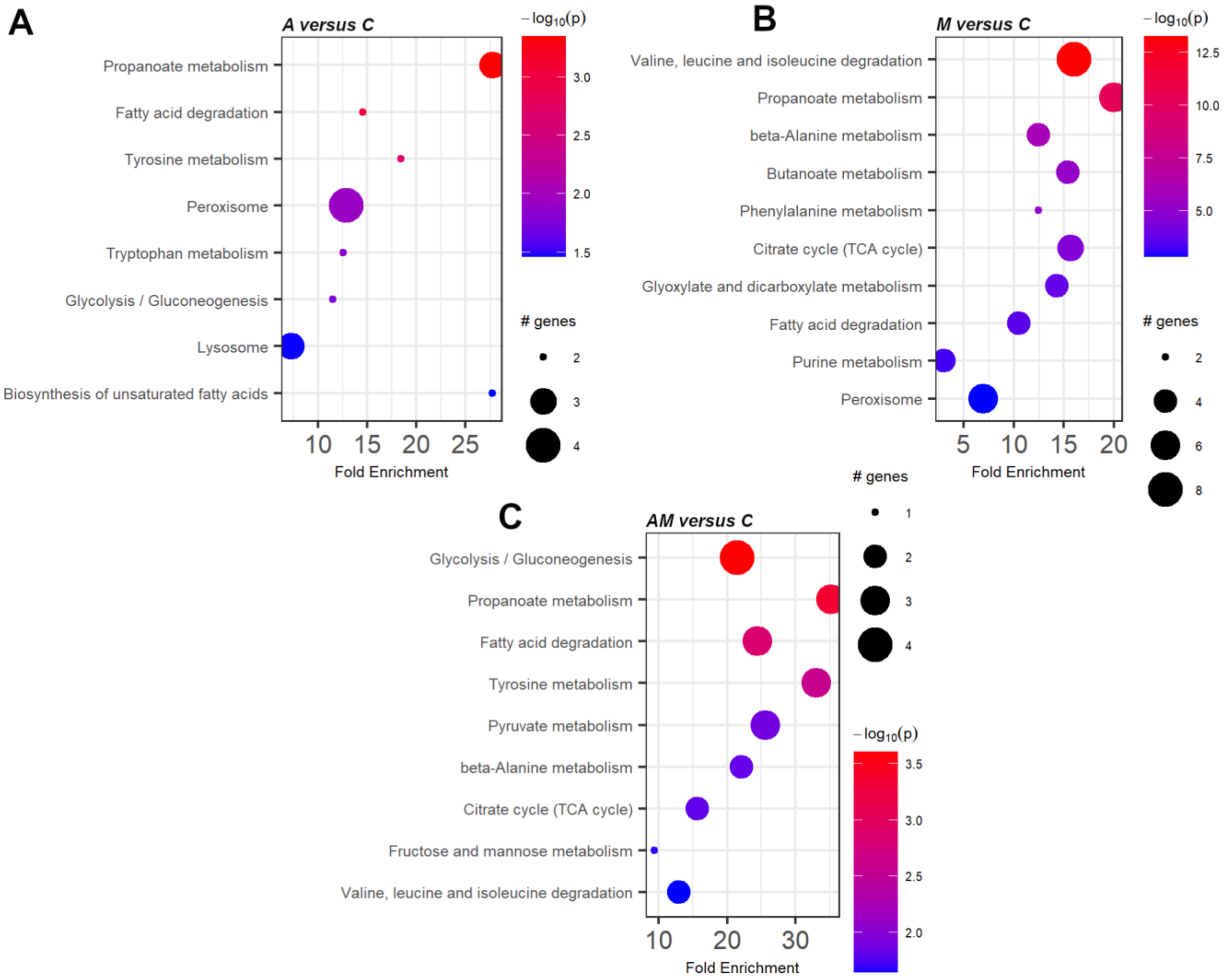


**Figure S2**. Enrichment chart from $log_2FC$ A/C (A), $log_2FC$ M/C (B) and $log_2FC$ AM/C (C) data, for the cytoplasmatic fraction from the liver of weaned piglets. The control group (C) were fed with the basal diet. The antioxidants experimental group were fed with the basal diet plus a mixture (1:1) of two byproducts (grapeseed and sea buckthorn meal) (A group).

Figure S3

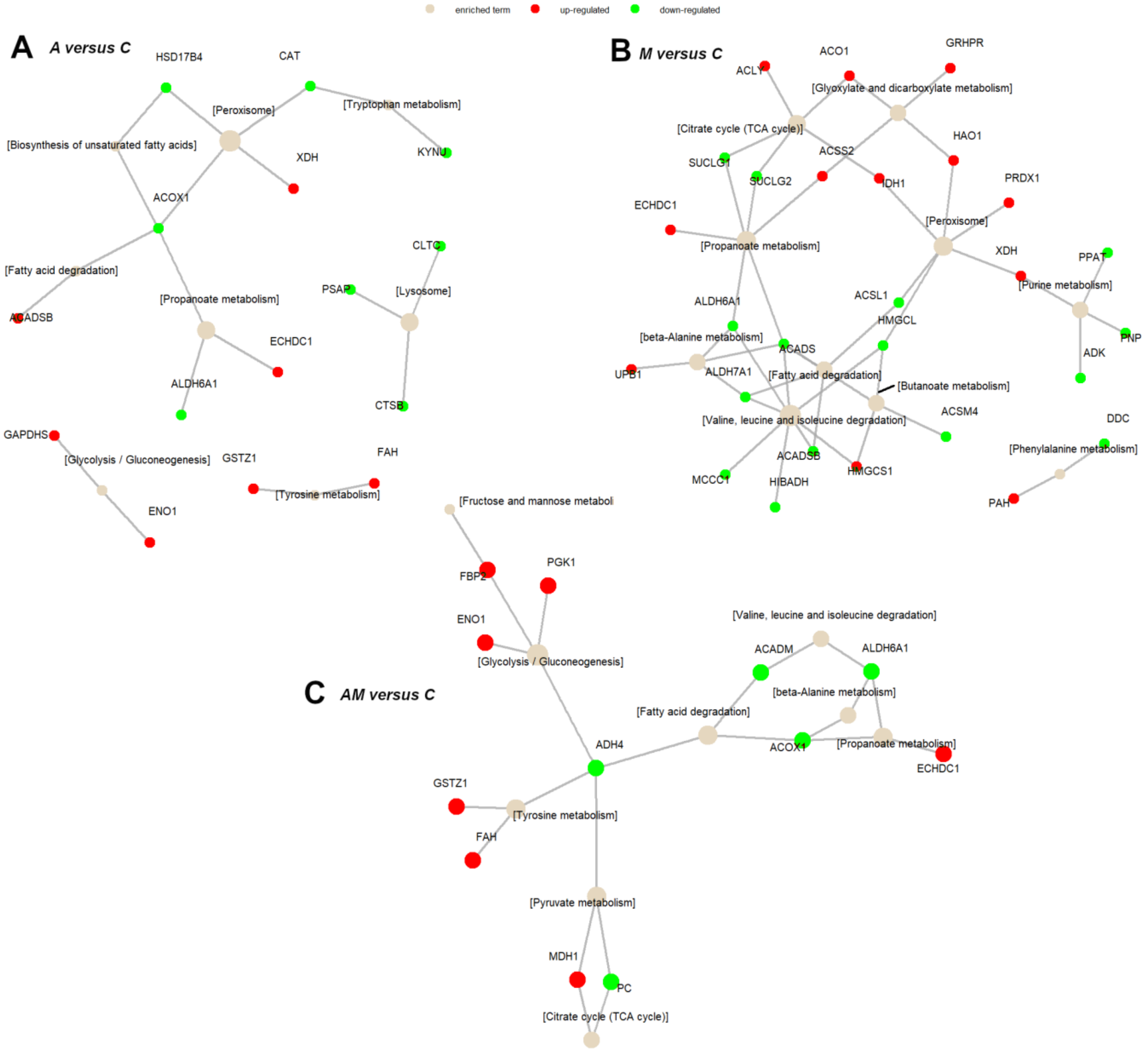


**Figure S3**. Term-gene graph for top 10 terms in liver cytoplasmatic fraction.

Figure S4

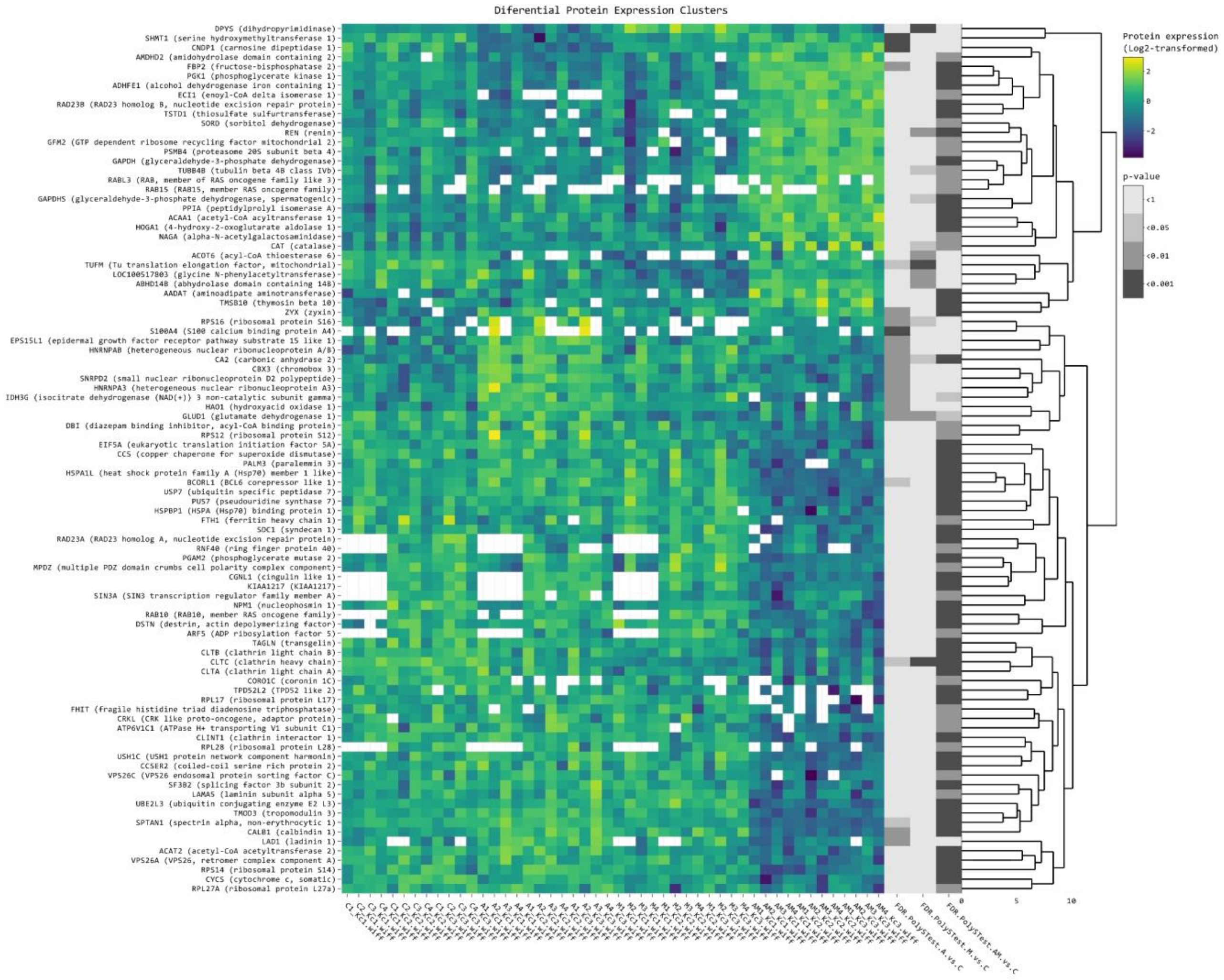


**Figure S4**. Clustered heatmap of the differentially expressed proteins in kideny cytoplasmatic fraction, filtered with log2FC threshold set to exclude interval (-0.5, 0.5) and FDR adjusted *p* value < 0.01. Yellow colour represents upregulation, while blue represents downregulation in A, M, and AM group compared to Control (C) group. From left to right, expression values (log2 transformed) for replicates (4 biological × 3 technical) are shown for the A, M, and AM groups, followed by significance values of the A versus C, M versus C, and AM versus C comparisons. The control group (C) was fed with a standard diet for starter piglets. The A group were fed with the basal diet plus a mixture of two antioxidant byproducts (grapeseed and sea buckthorn meal. The M group were fed with the basal diet artificially contaminated with two mycotoxins (AFB1 and OTA). The AM group represent the weaned piglets fed with the basal diet containing the mixture (1:1) of antioxidant byproducts and the two mycotoxins

Figure S5

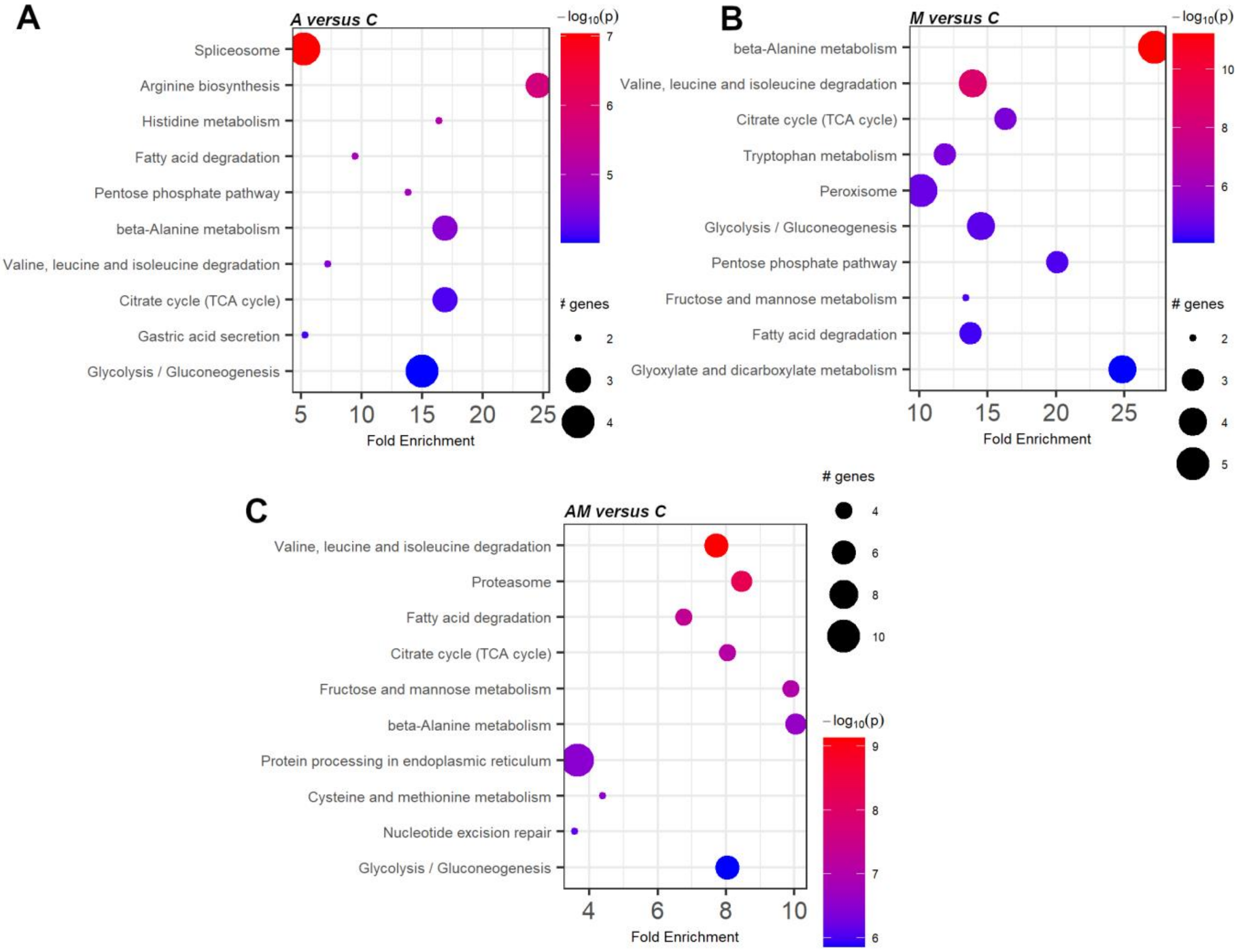


**Figure S5**. Enrichment chart for top 10 KEGG pathways in kidney cytoplasmatic fraction, sorted by lowest p value.

Figure S6

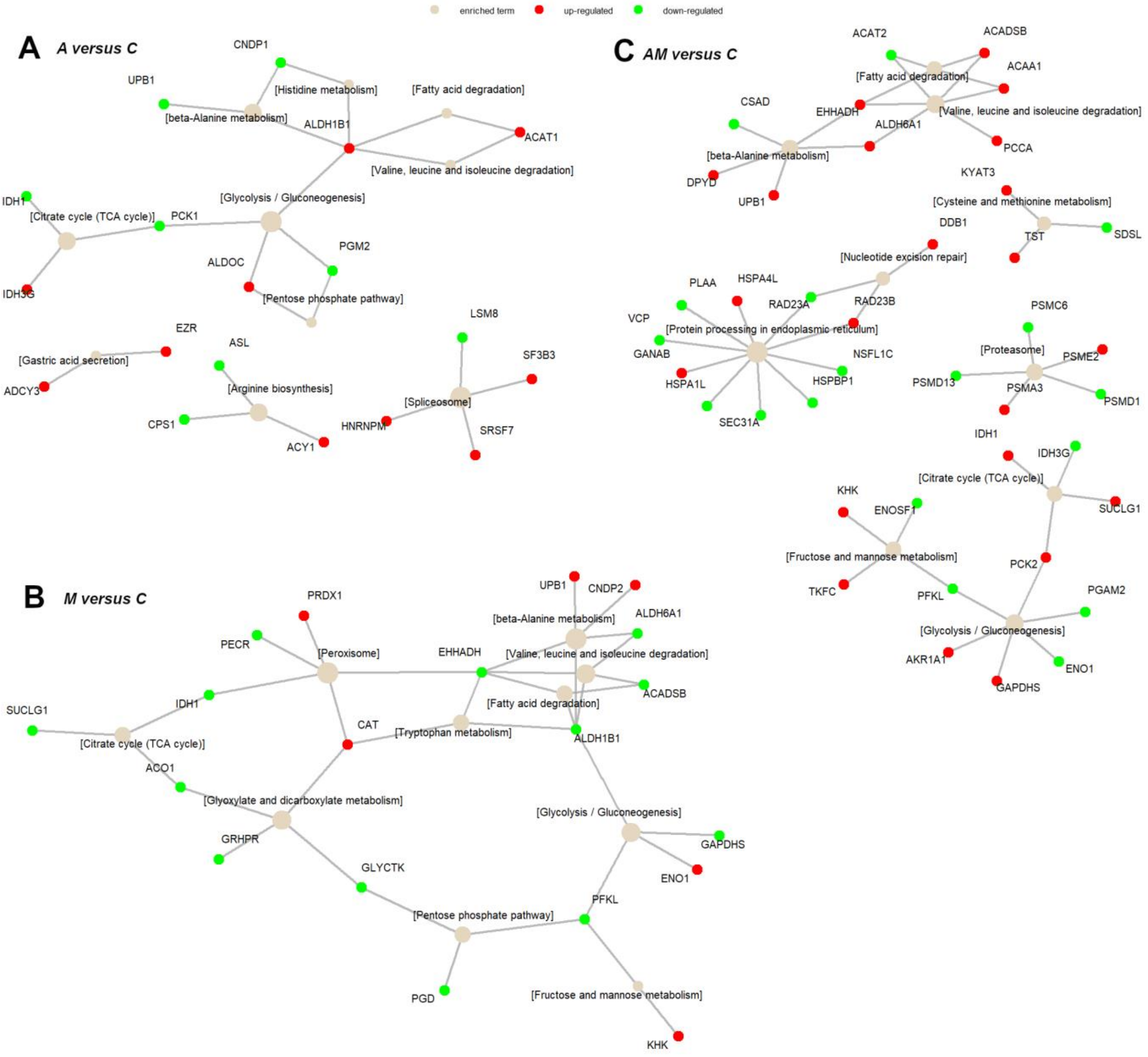


**Figure S6**. Term-gene graph for top 10 terms in kidney cytoplasmatic fraction.

Figure S7

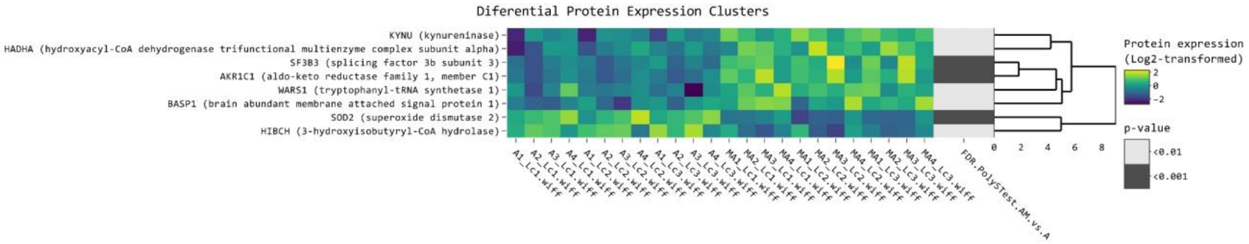


**Figure S7**. Mycotoxins impact in liver cytoplasmatic fraction. Clustered heatmap of the differentially expressed proteins (AM versus A), filtered with log2FC threshold set to exclude interval (-0.5, 0.5) and FDR adjusted *p* value < 0.01. Yellow colour represents upregulation, while blue represents downregulation in A group compared to AM group. From left to right, expression values (log2 transformed) for replicates (4 biological × 3 technical) are shown for the A group and for the AM group, followed by significance values of the comparison to A group. The A group were fed with the basal diet plus a mixture of two antioxidant byproducts (grapeseed and sea buckthorn meal. The AM group represent the weaned piglets fed with the basal diet containing the mixture (1:1) of antioxidant byproducts and the two mycotoxins.

Figure S8

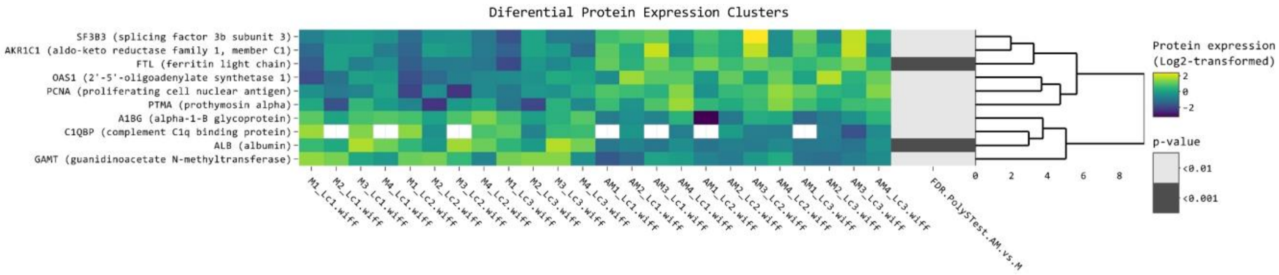


**Figure S8**. Antioxidants impact in liver cytoplasmatic fraction. Clustered heatmap of the differentially expressed proteins (AM versus M), filtered with log2FC threshold set to exclude interval (-0.5, 0.5) and FDR adjusted *p* value < 0.01. Yellow colour represents upregulation, while blue represents downregulation in A group compared to AM group. From left to right, expression values (log2 transformed) for replicates (4 biological × 3 technical) are shown for the M group and for the AM group, followed by significance values of the comparison to M group. The M group were fed with the basal diet artificially contaminated with two mycotoxins (AFB1 and OTA). The AM group represent the weaned piglets fed with the basal diet containing the mixture (1:1) of antioxidant byproducts and the two mycotoxins.

Figure S9

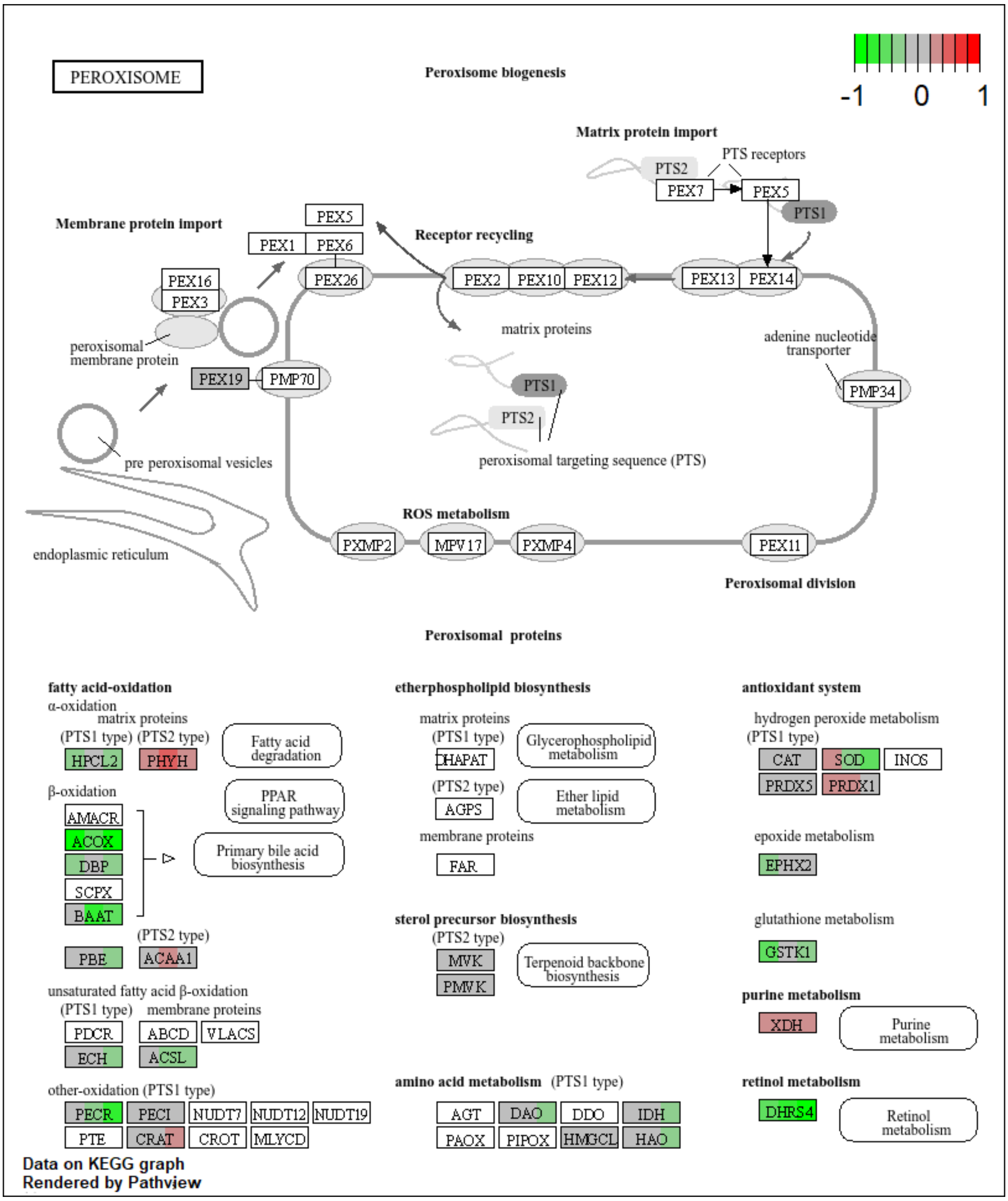


**Figure S9**. The KEGG peroxisome pathway for the cytoplasmatic fraction of the liver. The colour of the boxes represents the $\log_2$ fold change in the protein abundances, represented simultaneously for all three comparisons, on the left for A vs. C, in the middle for M vs. C and on the right for AM vs. C in the corresponding box for each protein.

Figure S10

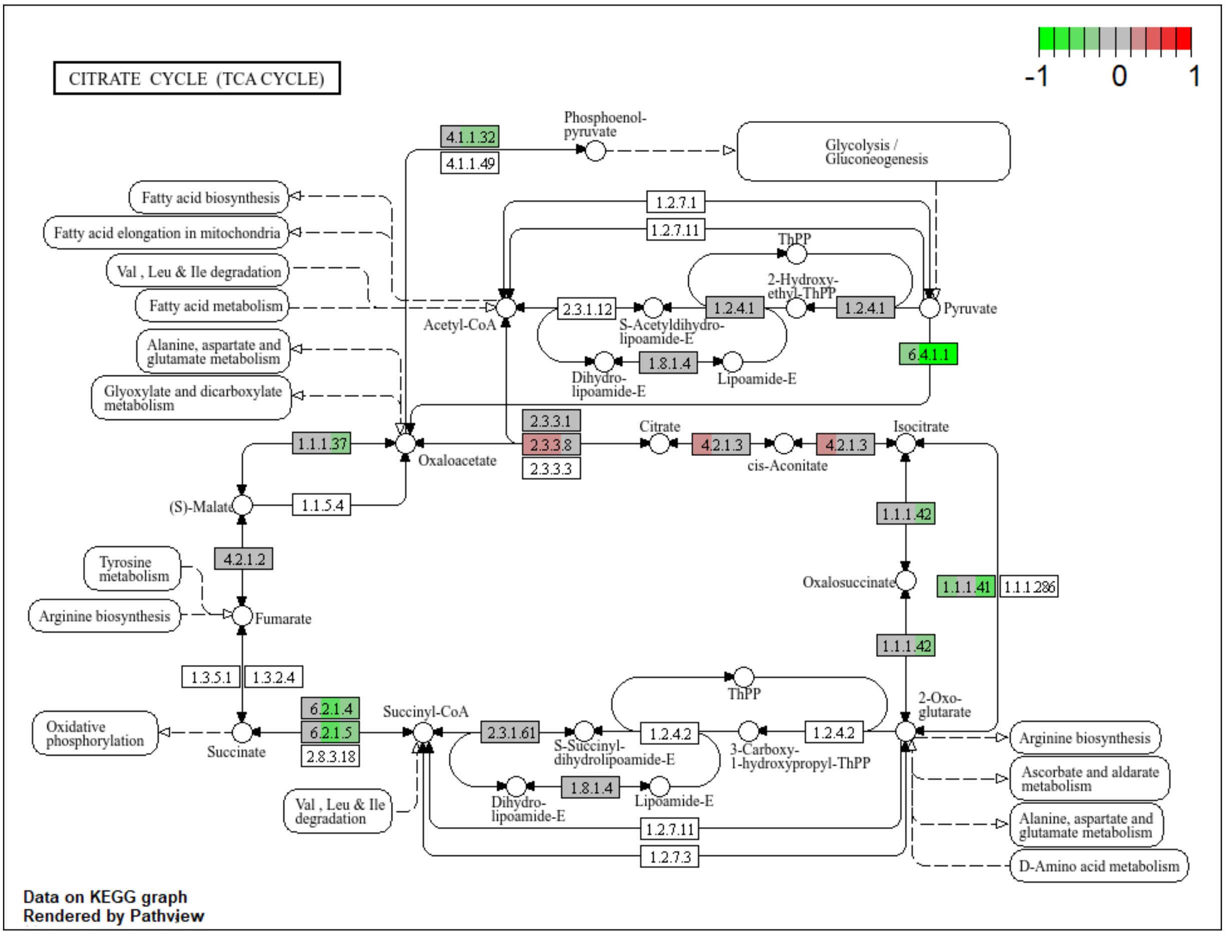


**Figure S10**. The KEGG TCA cycle for the cytoplasmatic fraction of the liver. The colour of the boxes represents the $log_2$ fold change in the protein abundances, represented simultaneously for all three comparisons, on the left for A vs. C, in the middle for M vs. C and on the right for AM vs. C in the corresponding box for each protein.

Figure S11

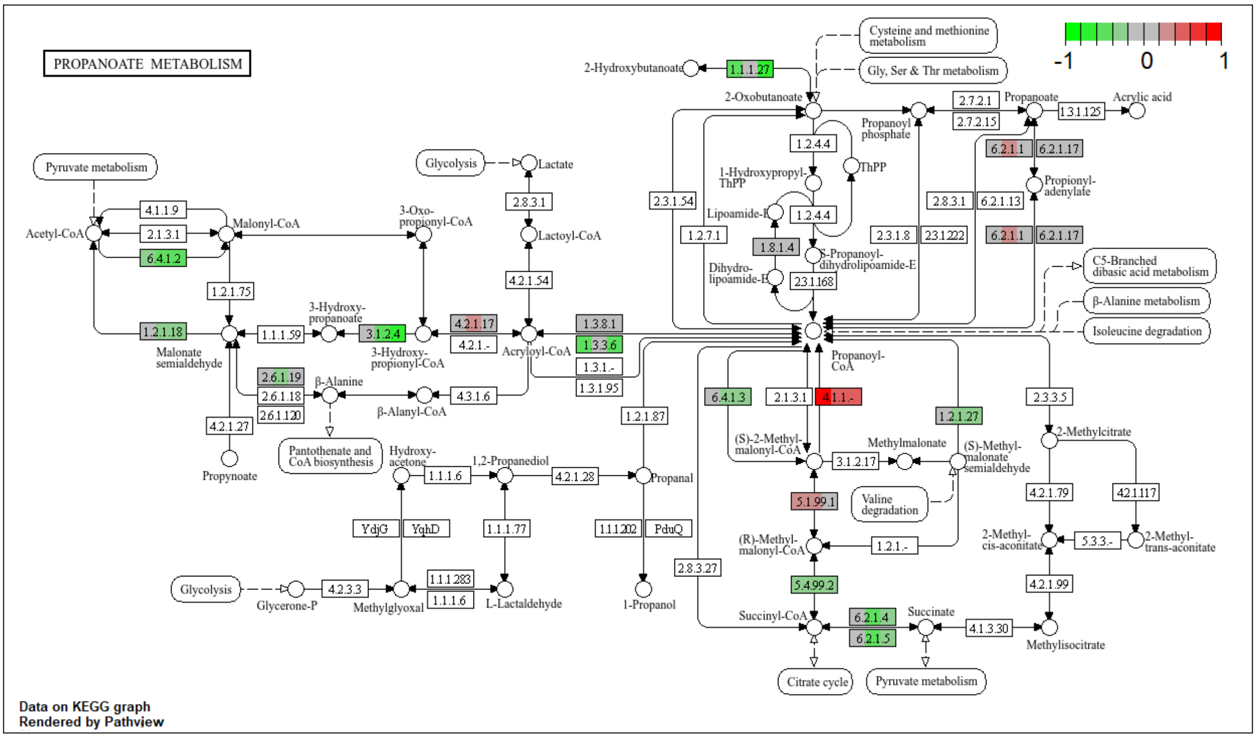


**Figure S11**. The KEGG propanoate metabolism for the cytoplasmatic fraction of the liver. The colour of the boxes represents the $\log_2$ fold change in the protein abundances, represented simultaneously for all three comparisons, on the left for A vs. C, in the middle for M vs. C and on the right for AM vs. C in the corresponding box for each protein.

Figure S12

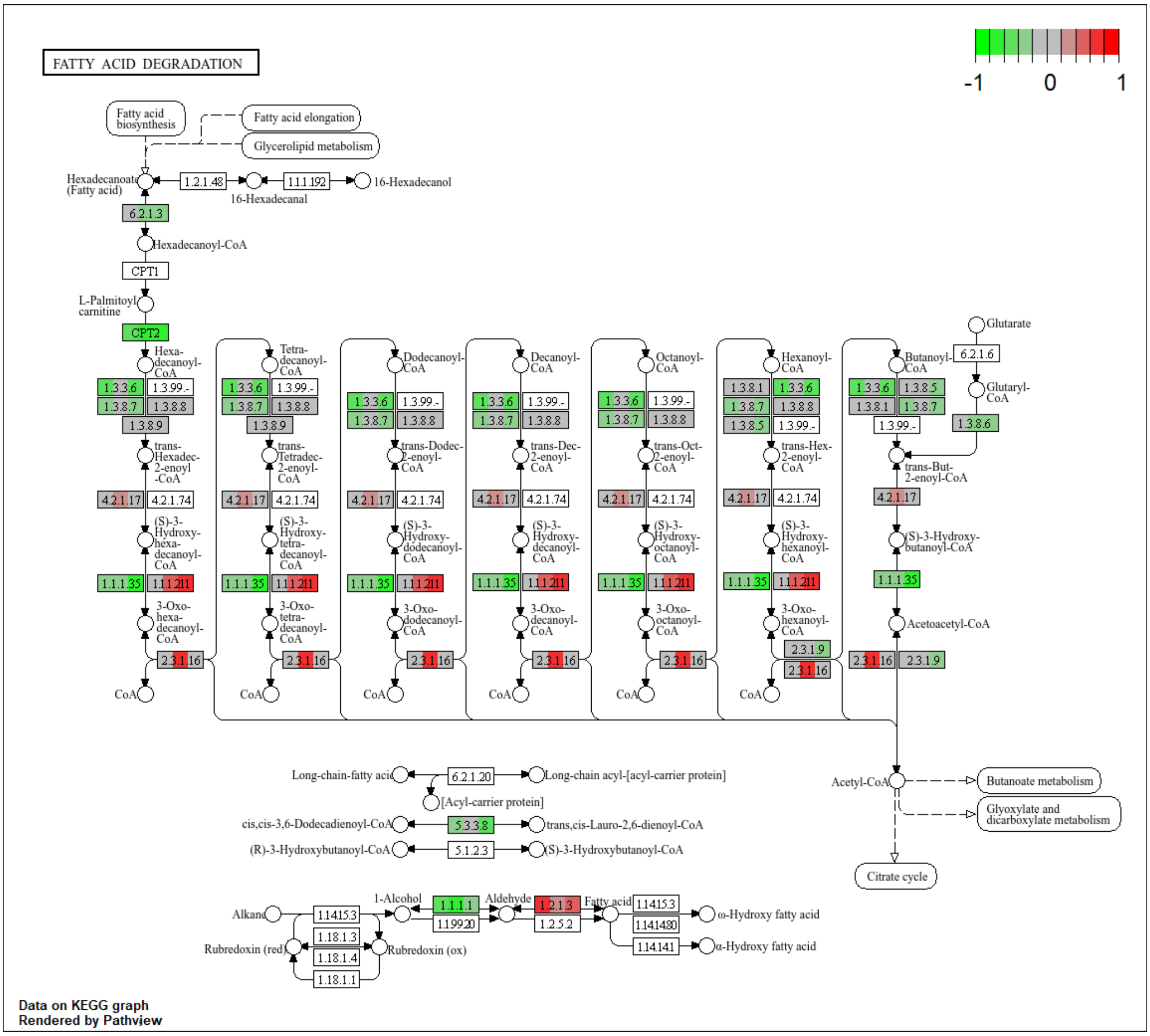


**Figure S12**. The KEGG fatty acid degradation for the cytoplasmatic fraction of the liver. The colour of the boxes represents the $\log_2$ fold change in the protein abundances, represented simultaneously for all three comparisons, on the left for A vs. C, in the middle for M vs. C and on the right for AM vs. C in the corresponding box for each protein.

Figure S13

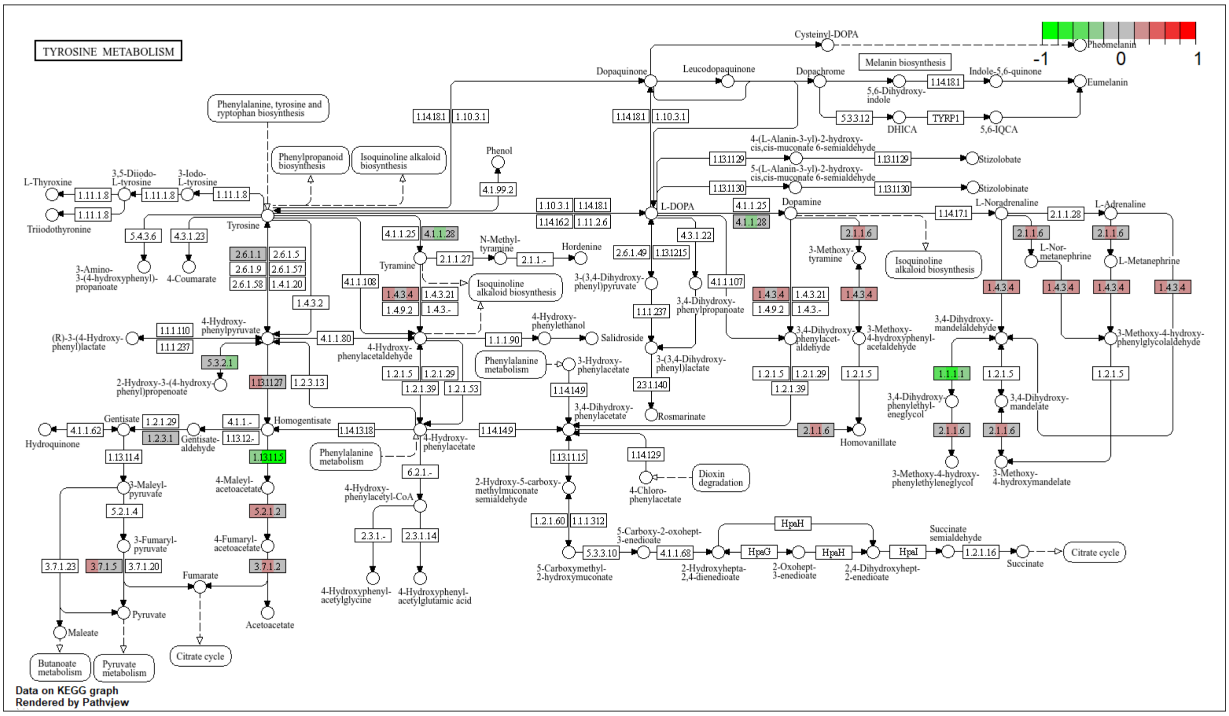


**Figure S13**. The KEGG tyrosine metabolism for the cytoplasmatic fraction of the liver. The colour of the boxes represents the $\log_2$ fold change in the protein abundances, represented simultaneously for all three comparisons, on the left for A vs. C, in the middle for M vs. C and on the right for AM vs. C in the corresponding box for each protein.

Figure S14

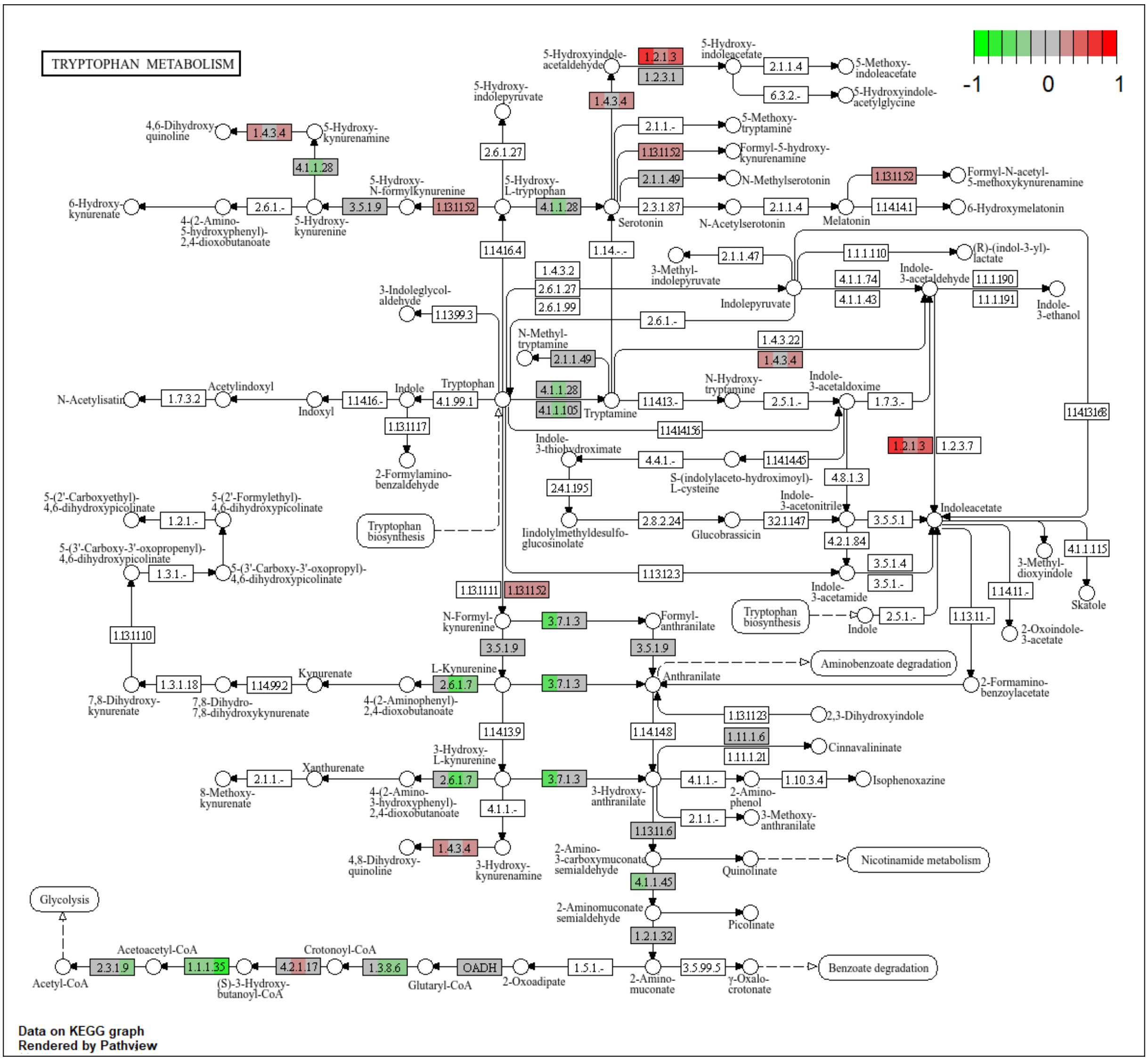


**Figure S14**. The KEGG tryptophan metabolism for the cytoplasmatic fraction of the liver. The colour of the boxes represents the $\log_2$ fold change in the protein abundances, represented simultaneously for all three comparisons, on the left for A vs. C, in the middle for M vs. C and on the right for AM vs. C in the corresponding box for each protein.

Figure S15

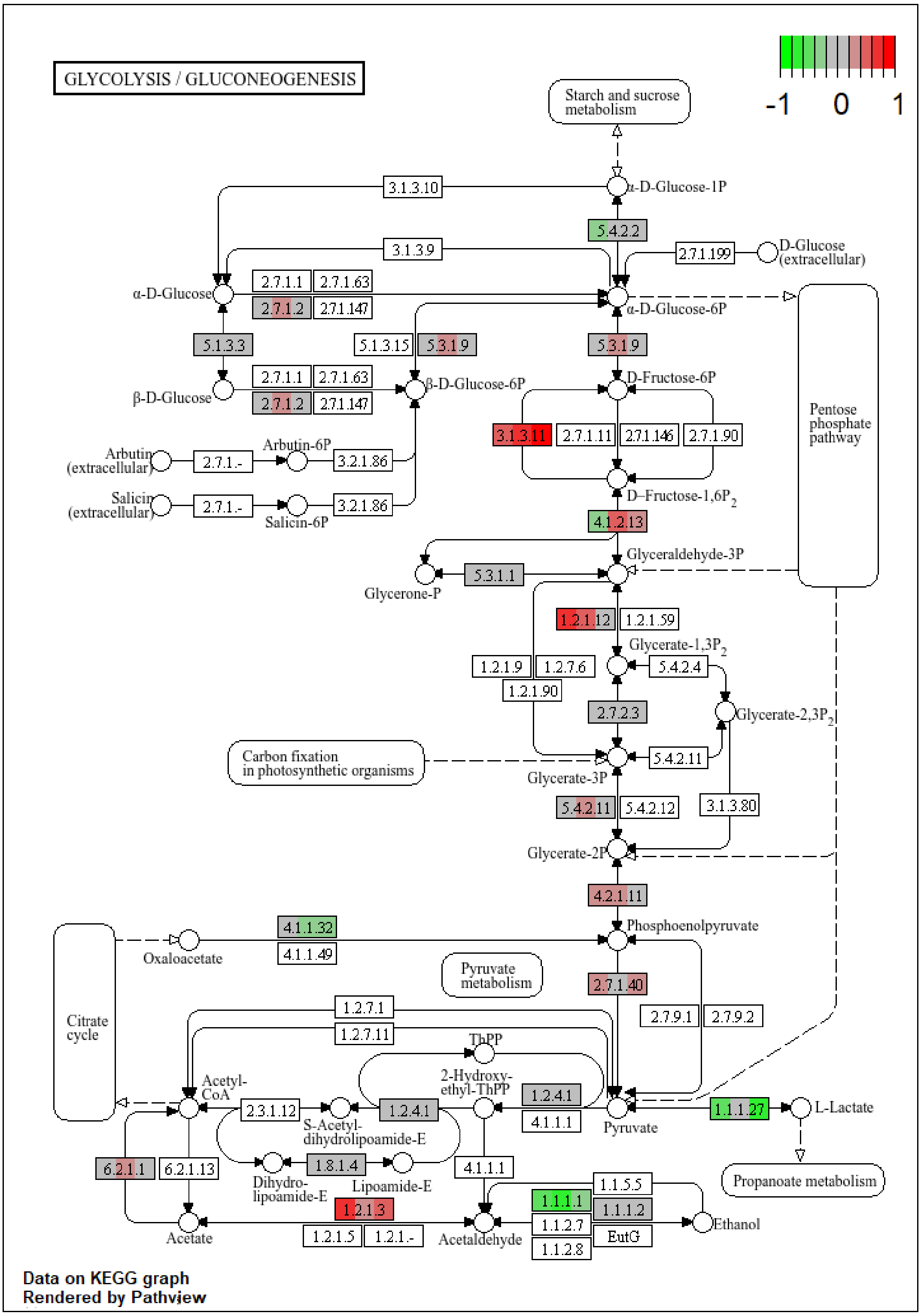


**Figure S15**. The KEGG glycolysis /gluconeogenesis pathway for the cytoplasmatic fraction of the liver. The colour of the boxes represents the $log_2$ fold change in the protein abundances, represented simultaneously for all three comparisons, on the left for A vs. C, in the middle for M vs. C and on the right for AM vs. C in the corresponding box for each protein.

Figure S16

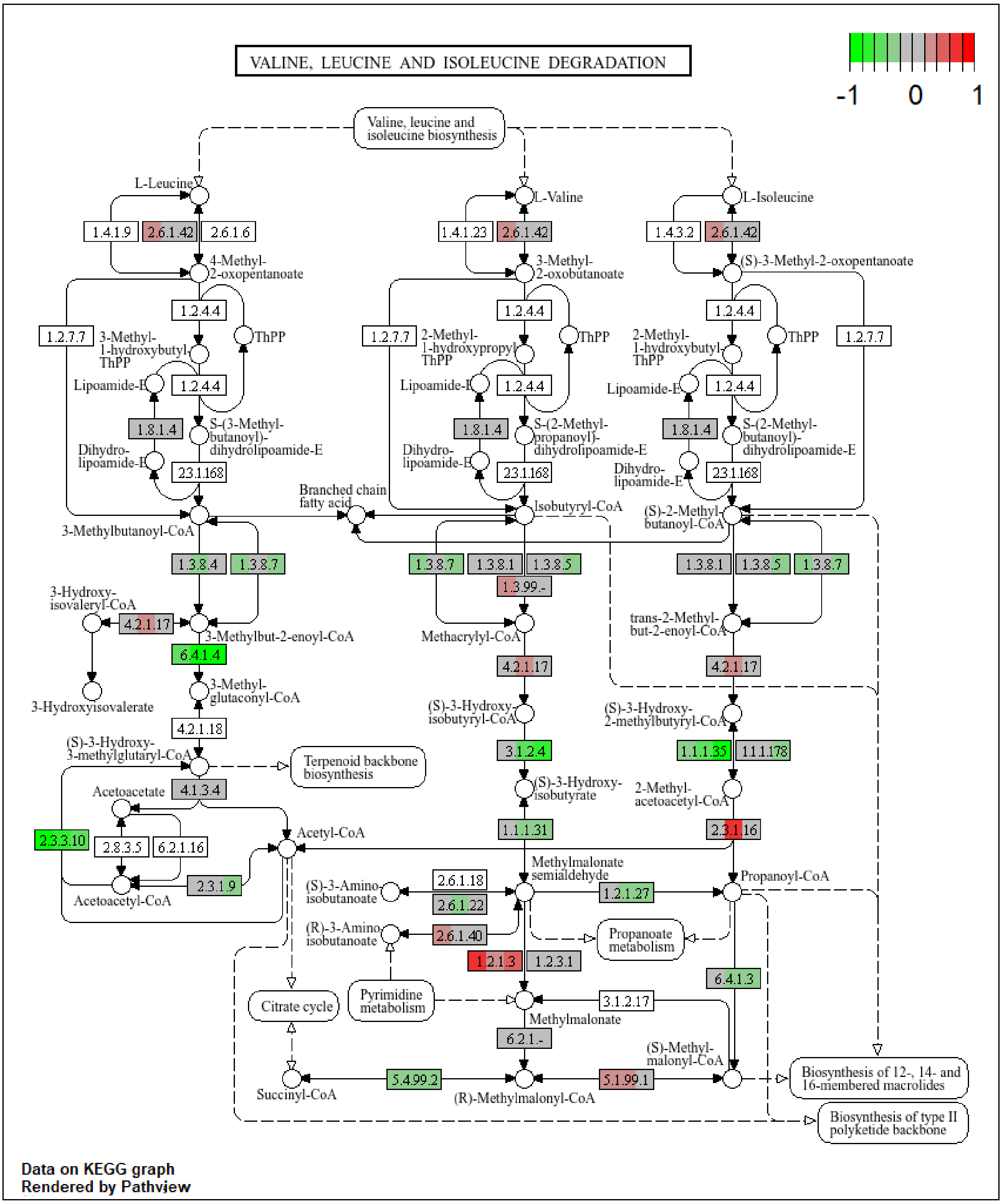


**Figure S16**. The KEGG valine, leucine and isoleucine degradation pathway for the cytoplasmatic fraction of the liver. The colour of the boxes represents the $\log_2$ fold change in the protein abundances, represented simultaneously for all three comparisons, on the left for A vs. C, in the middle for M vs. C and on the right for AM vs. C in the corresponding box for each protein.

Figure S17

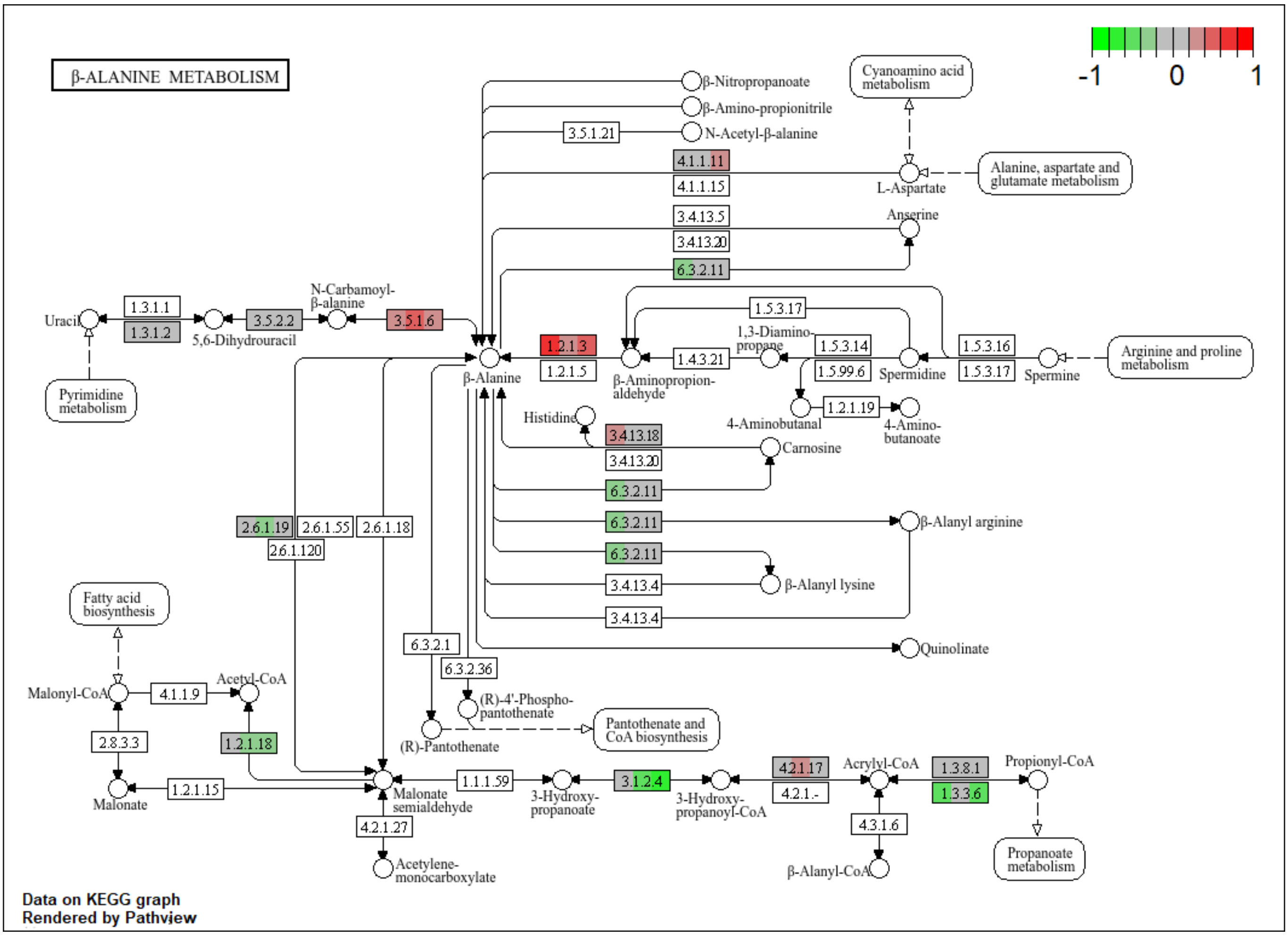


**Figure S17**. The KEGG beta-alanine pathway for the cytoplasmatic fraction of the liver. The colour of the boxes represents the $\log_2$ fold change in the protein abundances, represented simultaneously for all three comparisons, on the left for A vs. C, in the middle for M vs. C and on the right for AM vs. C in the corresponding box for each protein.

Figure S18

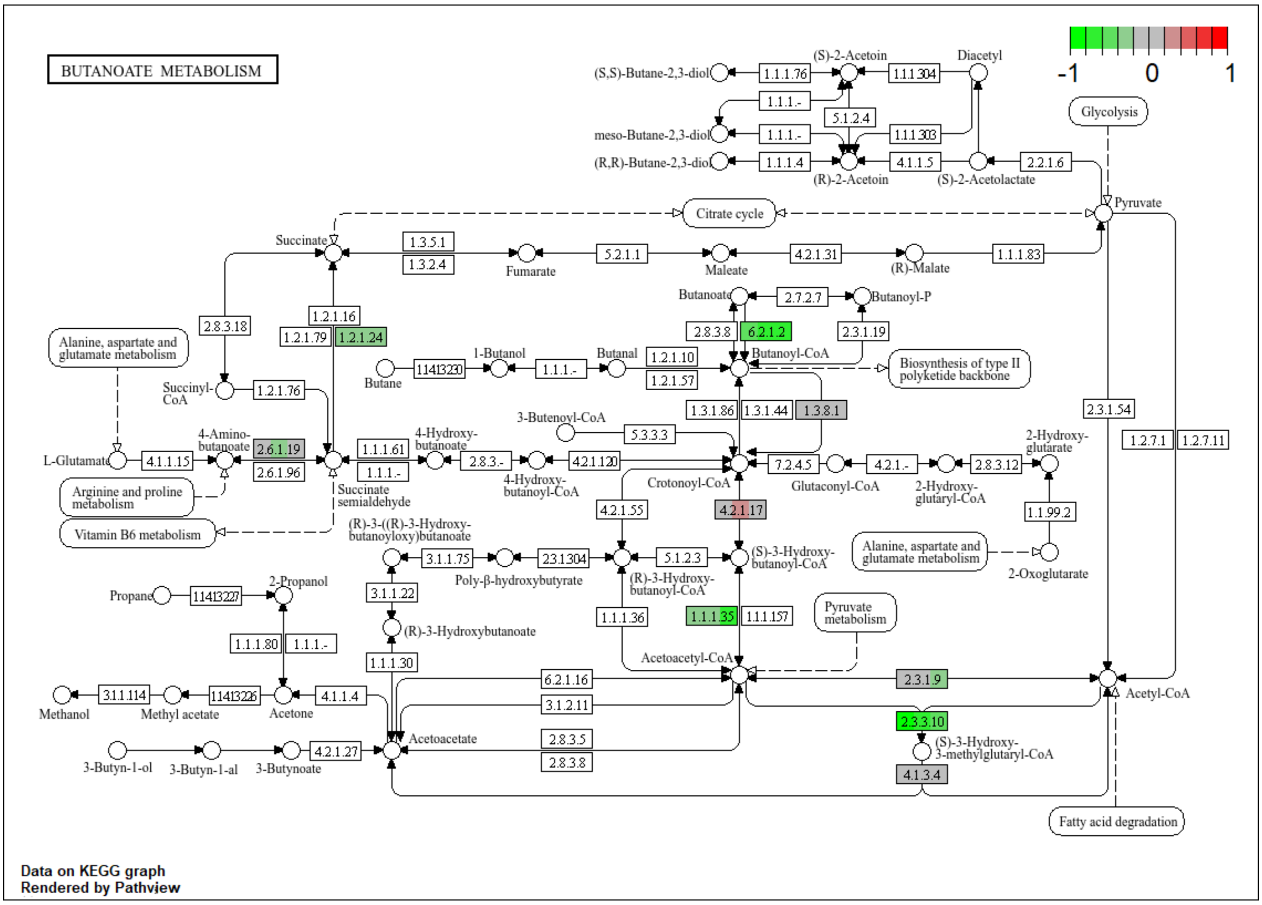


**Figure S18**. The KEGG butanoate metabolism for the cytoplasmatic fraction of the liver. The colour of the boxes represents the $\log_2$ fold change in the protein abundances, represented simultaneously for all three comparisons, on the left for A vs. C, in the middle for M vs. C and on the right for AM vs. C in the corresponding box for each protein.

Figure S19

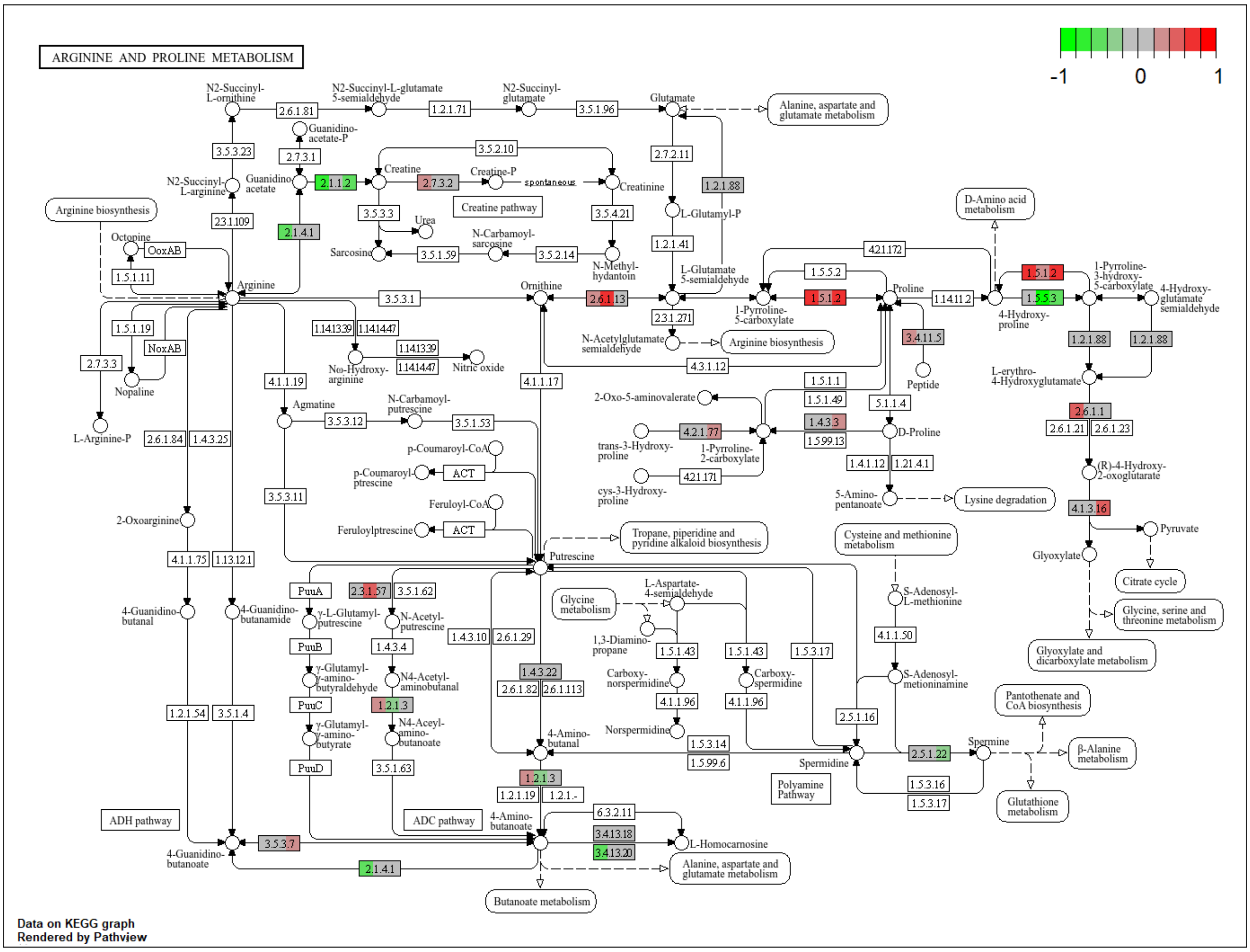


**Figure S19**. The KEGG arginine and proline metabolism for the cytoplasmatic fraction of the kidney. The colour of the boxes represents the $log_2$ fold change in the protein abundances, represented simultaneously for all three comparisons, on the left for A vs. C, in the middle for M vs. C and on the right for AM vs. C in the corresponding box for each protein.

Figure S20

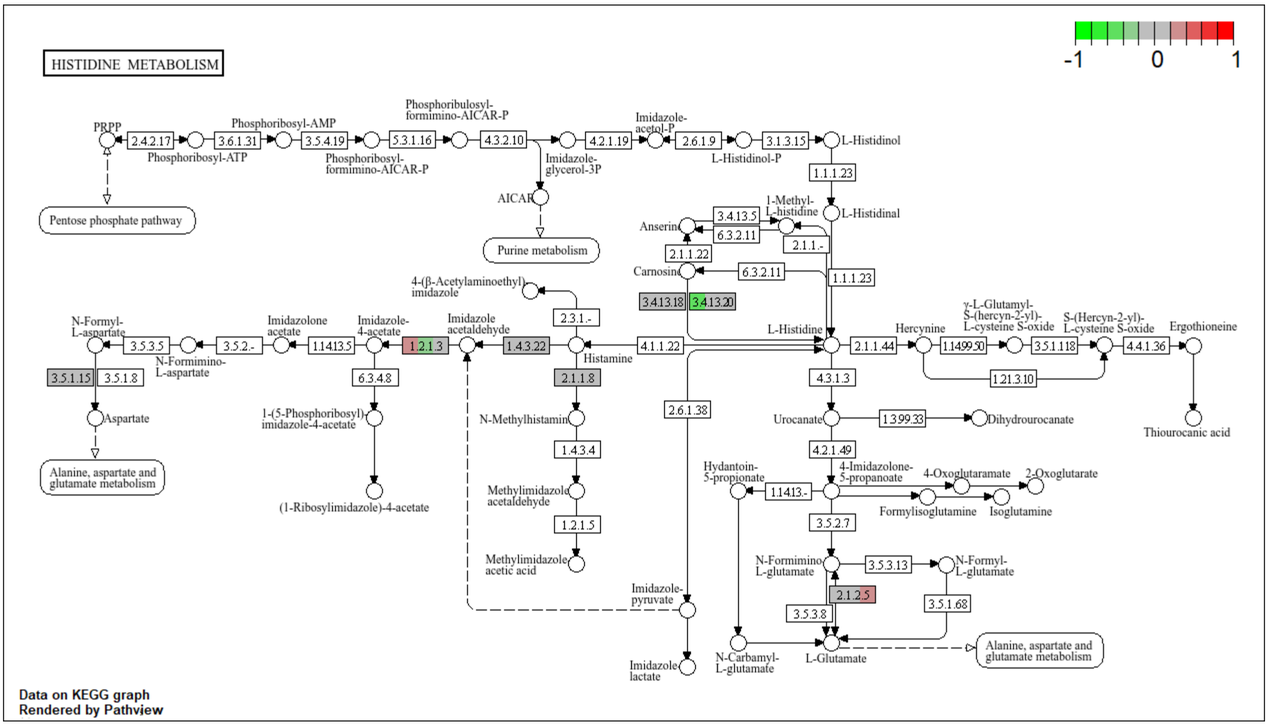


**Figure S20**. The KEGG histidine metabolism for the cytoplasmatic fraction of the kidney. The colour of the boxes represents the $\log_2$ fold change in the protein abundances, represented simultaneously for all three comparisons, on the left for A vs. C, in the middle for M vs. C and on the right for AM vs. C in the corresponding box for each protein.

Figure S21

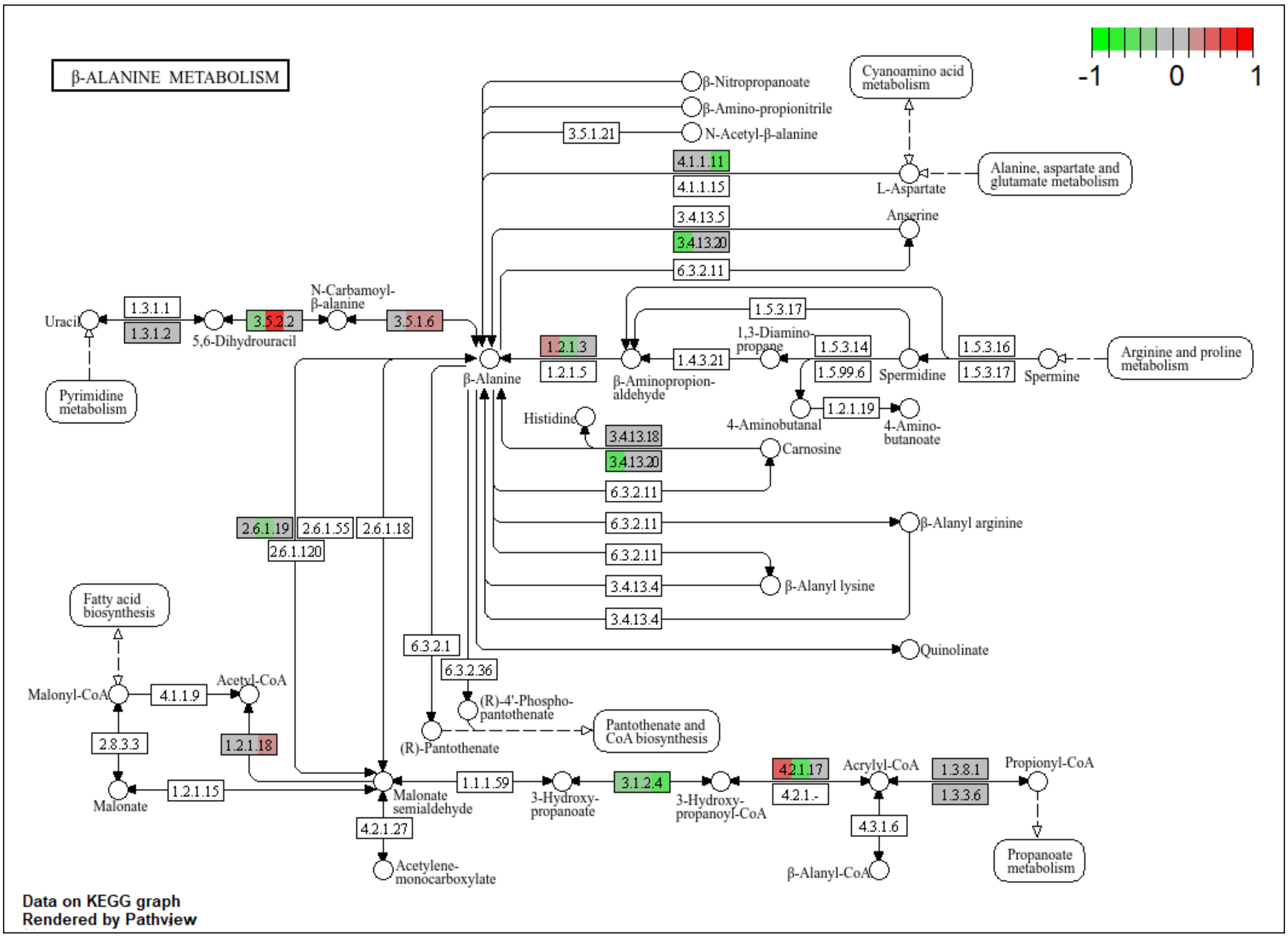


**Figure S21**. The KEGG beta-alanine metabolism for the cytoplasmatic fraction of the kidney. The colour of the boxes represents the $log_2$ fold change in the protein abundances, represented simultaneously for all three comparisons, on the left for A vs. C, in the middle for M vs. C and on the right for AM vs. C in the corresponding box for each protein.

Figure S22

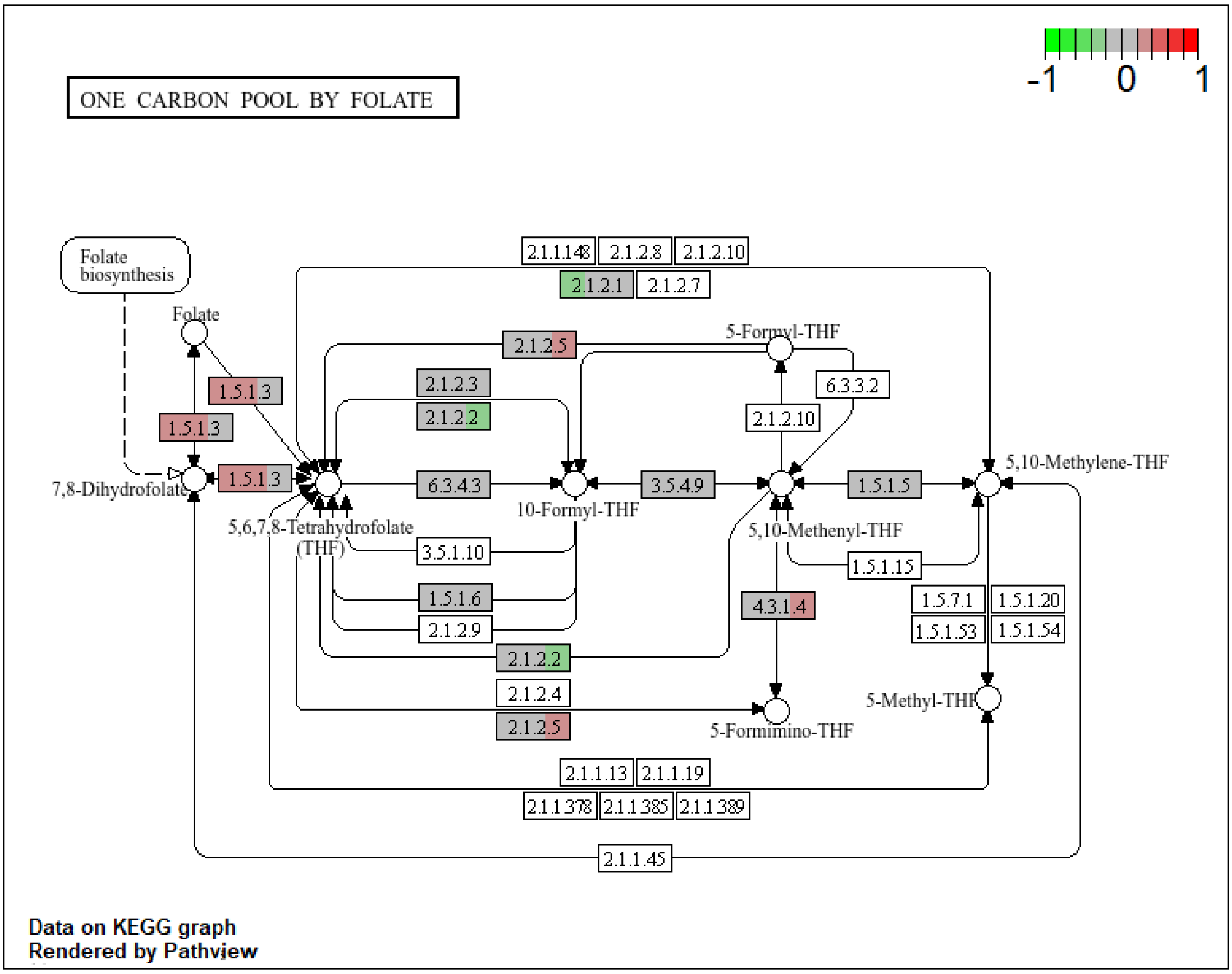


**Figure S22**. The KEGG one carbon pool by folate pathway for the cytoplasmatic fraction of the kidney. The colour of the boxes represents the $log_2$ fold change in the protein abundances, represented simultaneously for all three comparisons, on the left for A vs. C, in the middle for M vs. C and on the right for AM vs. C in the corresponding box for each protein.

Figure S23

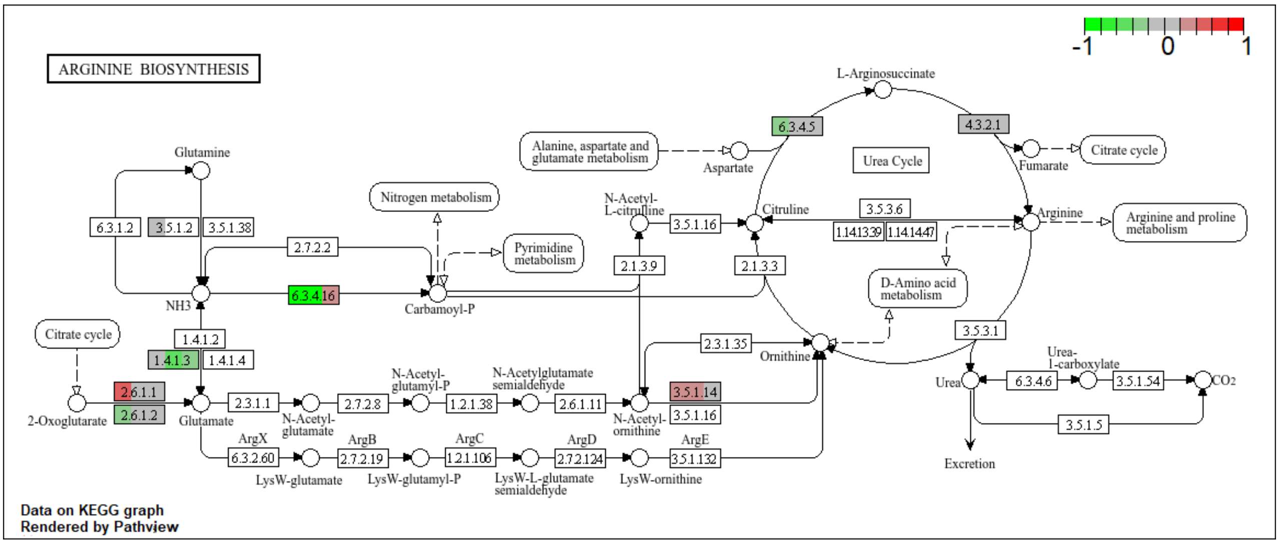


**Figure S23**. The KEGG arginine biosynthesis pathway for the cytoplasmatic fraction of the kidney. The colour of the boxes represents the $log_2$ fold change in the protein abundances, represented simultaneously for all three comparisons, on the left for A vs. C, in the middle for M vs. C and on the right for AM vs. C in the corresponding box for each protein.

Figure S24

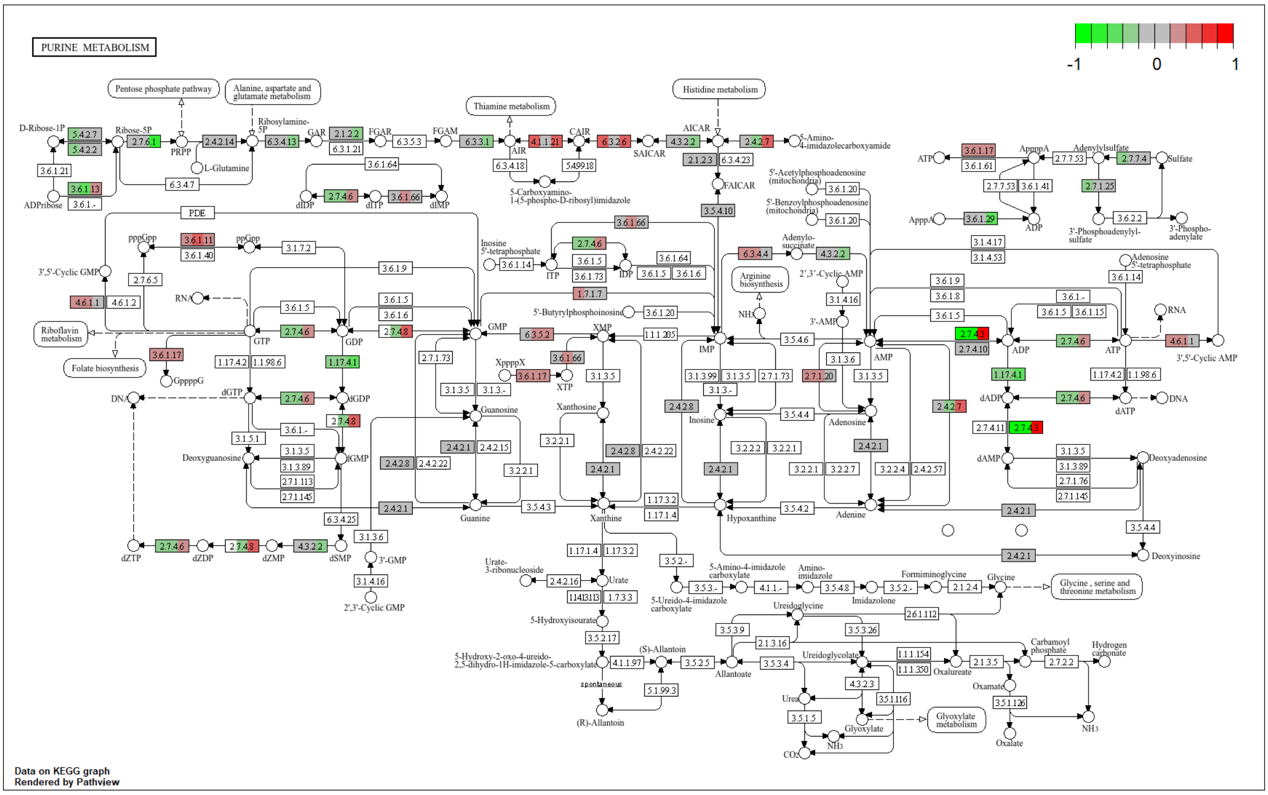


**Figure S24**. The KEGG purine metabolism for the cytoplasmatic fraction of the kidney. The colour of the boxes represents the $log_2$ fold change in the protein abundances, represented simultaneously for all three comparisons, on the left for A vs. C, in the middle for M vs. C and on the right for AM vs. C in the corresponding box for each protein.

Figure S25

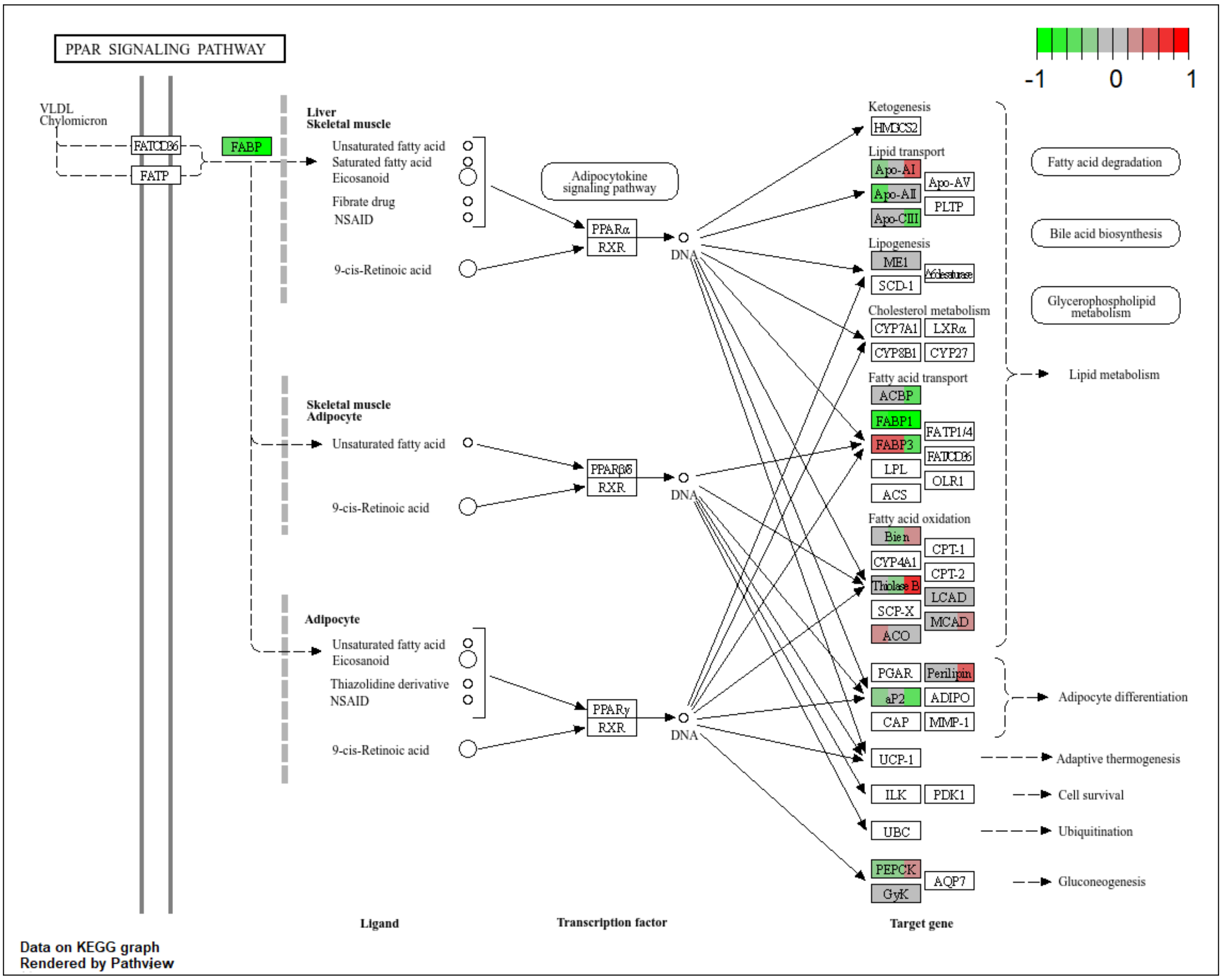


**Figure S25**. The KEGG PPAR signalling pathway for the cytoplasmatic fraction of the kidney. The colour of the boxes represents the $log_2$ fold change in the protein abundances, represented simultaneously for all three comparisons, on the left for A vs. C, in the middle for M vs. C and on the right for AM vs. C in the corresponding box for each protein.

Figure S26

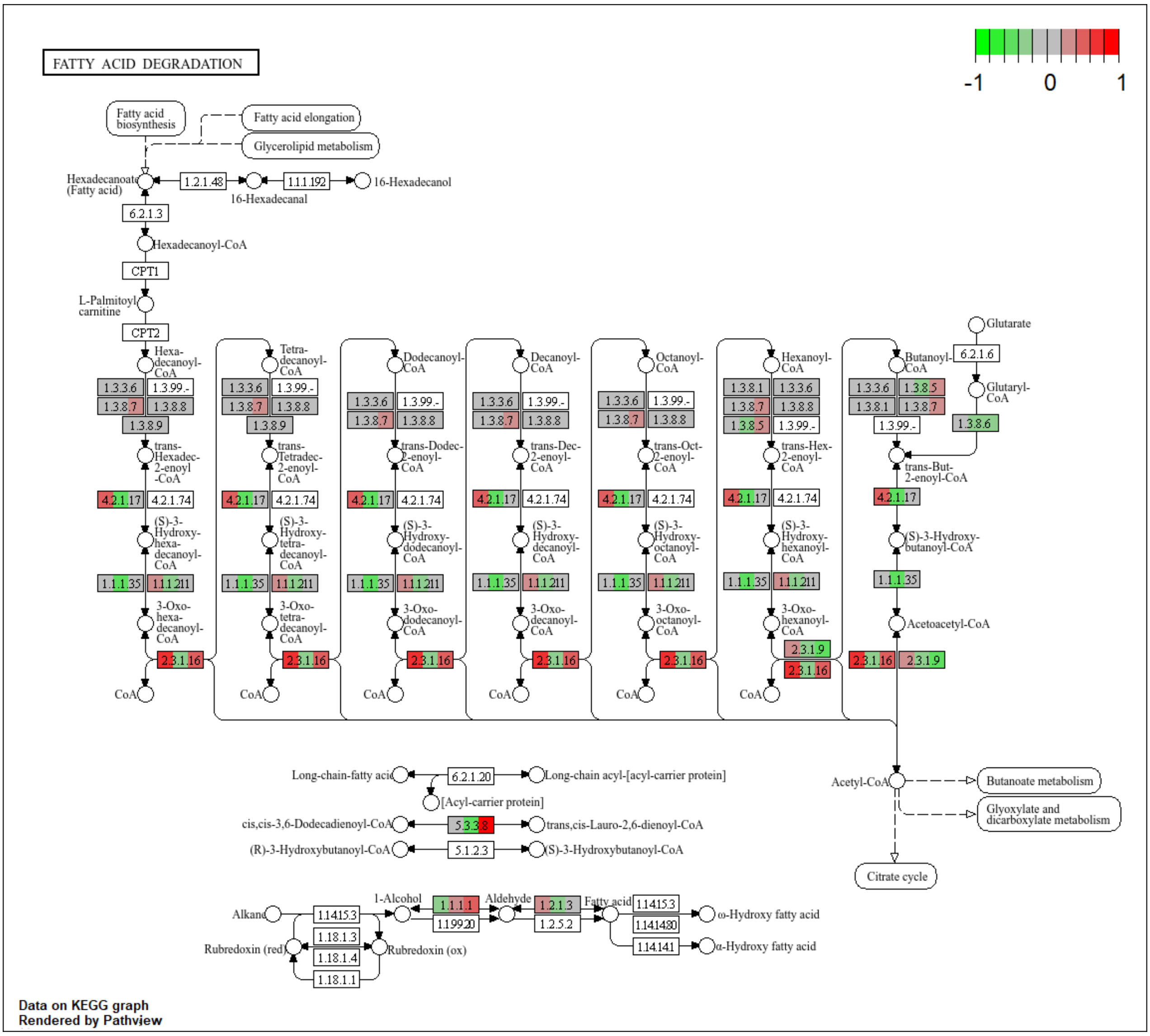


**Figure S26**. The KEGG fatty acid degradation pathway for the cytoplasmatic fraction of the kidney. The colour of the boxes represents the $log_2$ fold change in the protein abundances, represented simultaneously for all three comparisons, on the left for A vs. C, in the middle for M vs. C and on the right for AM vs. C in the corresponding box for each protein.

Figure S27

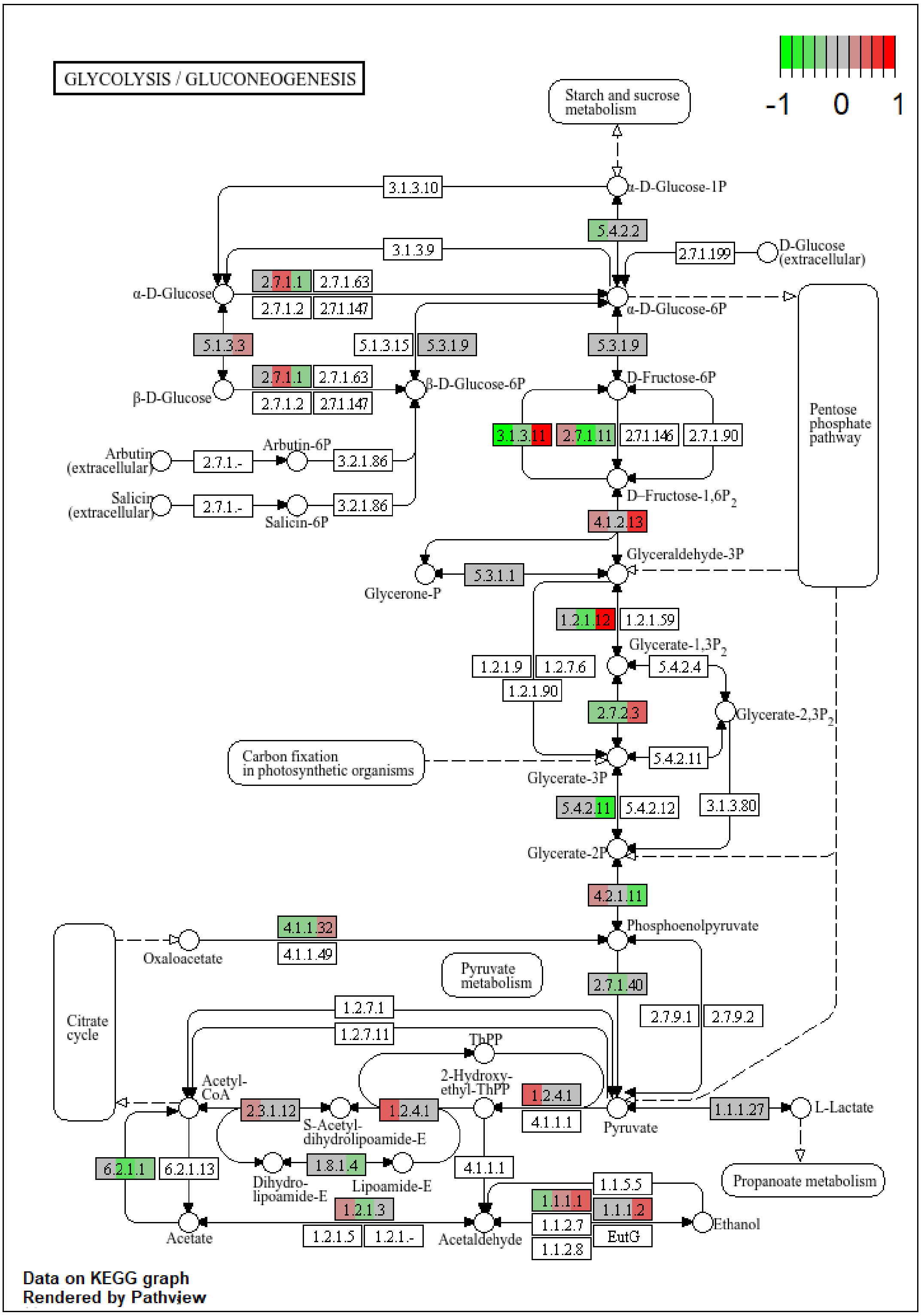


**Figure S27**. The KEGG glycolysis /gluconeogenesis pathway for the cytoplasmatic fraction of the kidney. The colour of the boxes represents the $log_2$ fold change in the protein abundances, represented simultaneously for all three comparisons, on the left for A vs. C, in the middle for M vs. C and on the right for AM vs. C in the corresponding box for each protein.

Figure S28

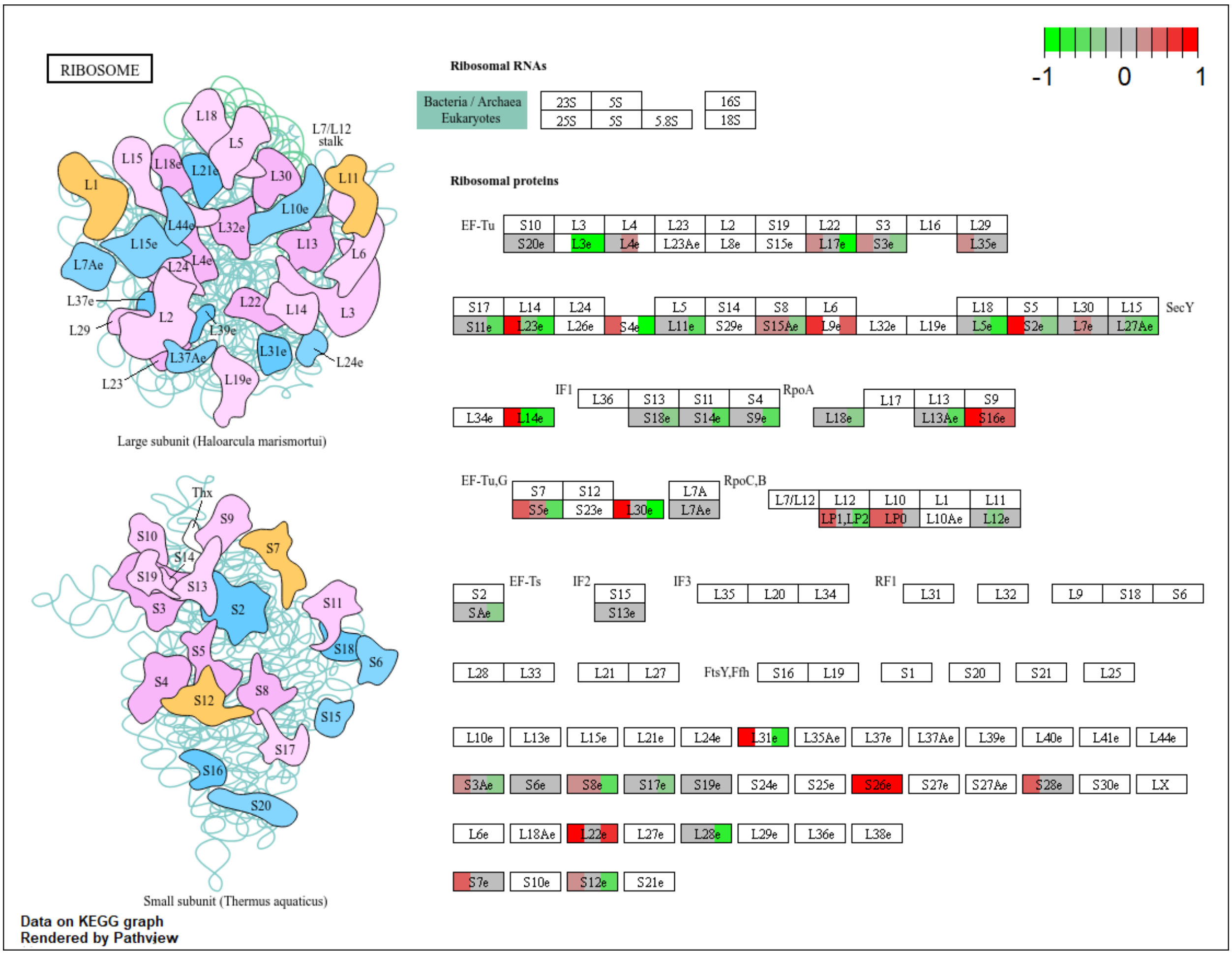


**Figure S28**. The KEGG ribosome pathway for the cytoplasmatic fraction of the kidney. The colour of the boxes represents the $log_2$ fold change in the protein abundances, represented simultaneously for all three comparisons, on the left for A vs. C, in the middle for M vs. C and on the right for AM vs. C in the corresponding box for each protein.

Figure S29

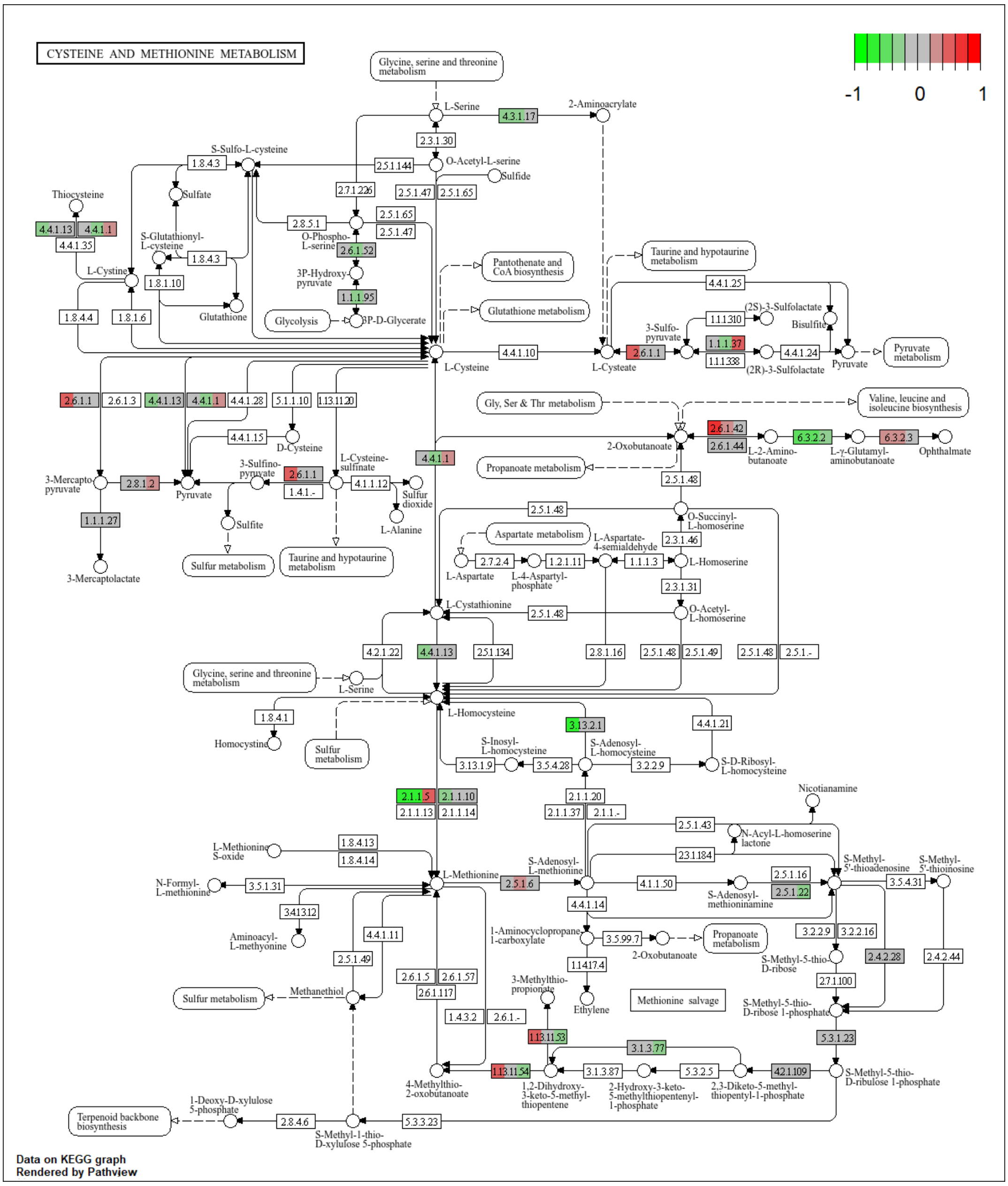

**Figure S29**. The KEGG cysteine and methionine metabolism for the cytoplasmatic fraction of the kidney. The colour of the boxes represents the $\log_2$ fold change in the protein abundances, represented simultaneously for all three comparisons, on the left for A vs. C, in the middle for M vs. C and on the right for AM vs. C in the corresponding box for each protein.

Figure S30

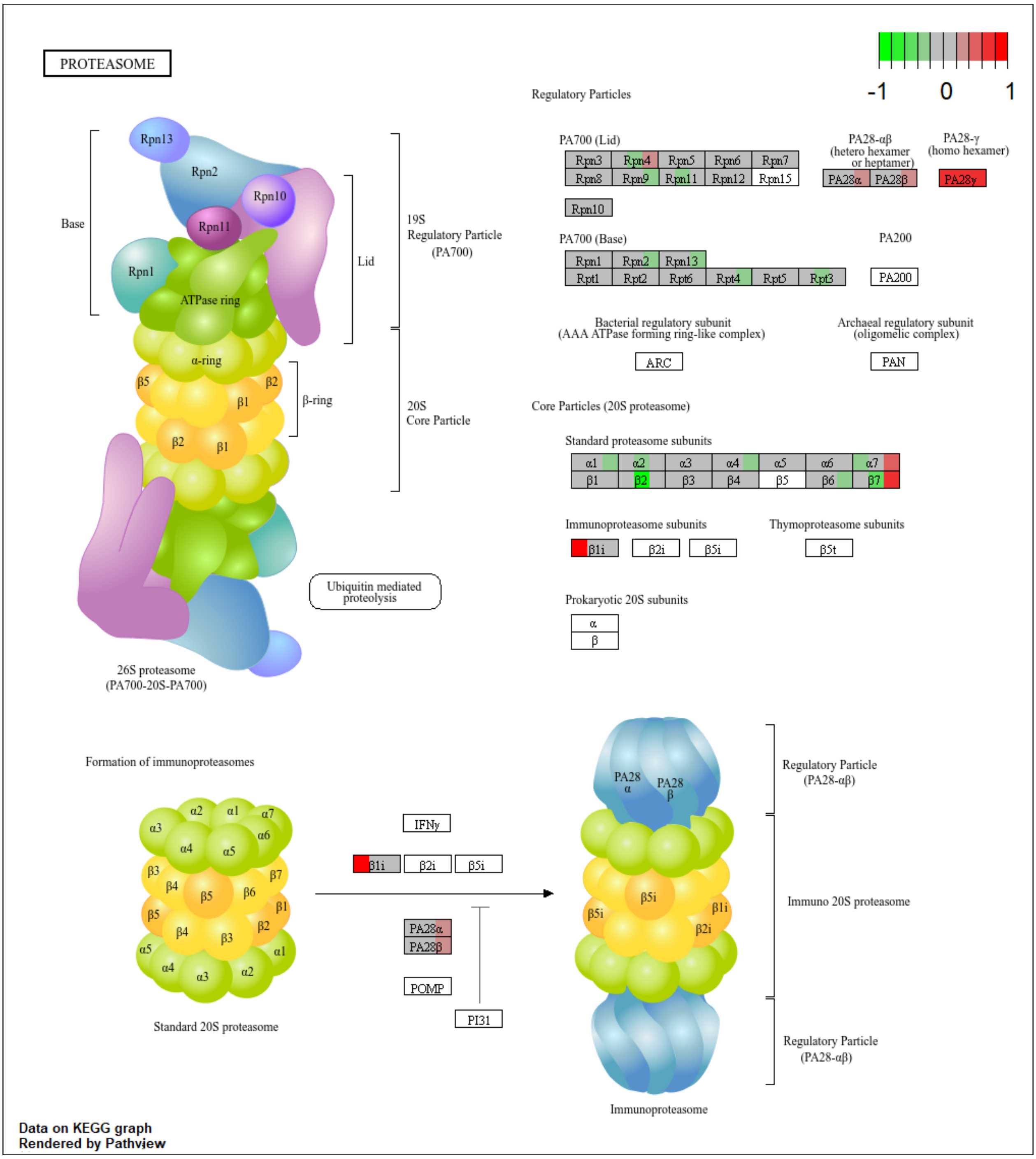


**Figure S30**. The KEGG proteasome pathway for the cytoplasmatic fraction of the kidney. The colour of the boxes represents the $log_2$ fold change in the protein abundances, represented simultaneously for all three comparisons, on the left for A vs. C, in the middle for M vs. C and on the right for AM vs. C in the corresponding box for each protein.
